# Cryo-EM Reveals How Cardiomyopathy Therapeutic Drugs Modulate the Myosin Motors of the Heart

**DOI:** 10.1101/2025.10.29.685122

**Authors:** Arun Kumar Somavarapu, Jinghua Ge, Christopher M. Yengo, Roger Craig, Raul Padron

## Abstract

Genetic mutations in myosin, the motor protein that powers the heartbeat, are linked to inherited hypertrophic and dilated cardiomyopathies. Mavacamten and omecamtiv mecarbil are therapeutic, myosin-targeted drugs designed to treat these myopathies, but their mechanism of action has remained unclear. Using single-particle cryo-EM, we determined near-atomic resolution structures of wild-type, mavacamten-bound, and omecamtiv mecarbil-bound myosin molecules. Across all conditions, two conformations of myosin were observed. We show how mavacamten stabilizes one conformation by reinforcing key electrostatic interfaces in the molecule, whereas omecamtiv mecarbil weakens these interfaces, favoring the second structure. This remodeling elucidates previously unclear allosteric mechanisms through which these drugs either inhibit or enhance myosin activity, countering the deleterious impacts of disease. These findings reveal how drugs modulate myosin structure to control cardiac contractility.

## Introduction

Beta-cardiac myosin is the two-headed molecular motor responsible for converting chemical energy from ATP hydrolysis into mechanical force, the fundamental basis of heart muscle contraction (*1, 2*). Each β-cardiac myosin molecule functions as a hexamer composed of two heavy chains (MHC), two essential light chains (ELC), and two regulatory light chains (RLC). Each heavy chain is organized into distinct regions with specific roles in force production and filament assembly. The N-terminal half of each heavy chain constitutes the motor domain, which holds the catalytic site for ATP hydrolysis and the actin-binding interface. This domain is further segmented into four subdomains: the N-terminal domain, the upper and lower 50 kDa domains (U50 and L50), and the converter domain. Immediately C-terminal to the motor domain is the light chain binding domain (LCD), which is associated with one ELC and one RLC per head, contributing to structural stability and regulatory control(*3, 4*). Following the LCD, each heavy chain extends into a long α-helix that dimerizes with the other through coiled-coil interactions, forming the myosin tail. The tail is divided into two functional regions: the proximal subfragment-2 (S2) and the distal light meromyosin (LMM). S2 plays a critical role in head-tail interactions (see below) while LMM drives self-association of myosin molecules to form the thick filament backbone(*5, 6*).

During the muscle contractile cycle, this two-headed molecular motor operates through two major structural states: an open, disordered-relaxed (DRX) state, where the two heads are separated from each other, ready to interact with actin; and a closed, super-relaxed (SRX) state (*7, 8*), in which the heads interact with each other, conserving energy (Fig. 1). The interacting heads (blocked and free, BH and FH) fold back onto their coiled-coil S2 tail, forming the interacting-heads motif (IHM), a conformation that dramatically reduces ATPase activity and actin binding(*9–12*). Cryo-EM studies reveal how BH-FH and BH-S2 interfaces stabilize the IHM(*11, 13*). The IHM is a conserved, energy-saving mechanism across muscle types (skeletal, cardiac, and smooth), and is also observed in non-muscle cells and even in primitive unicellular organisms(*14–16*).

**Fig 1.**
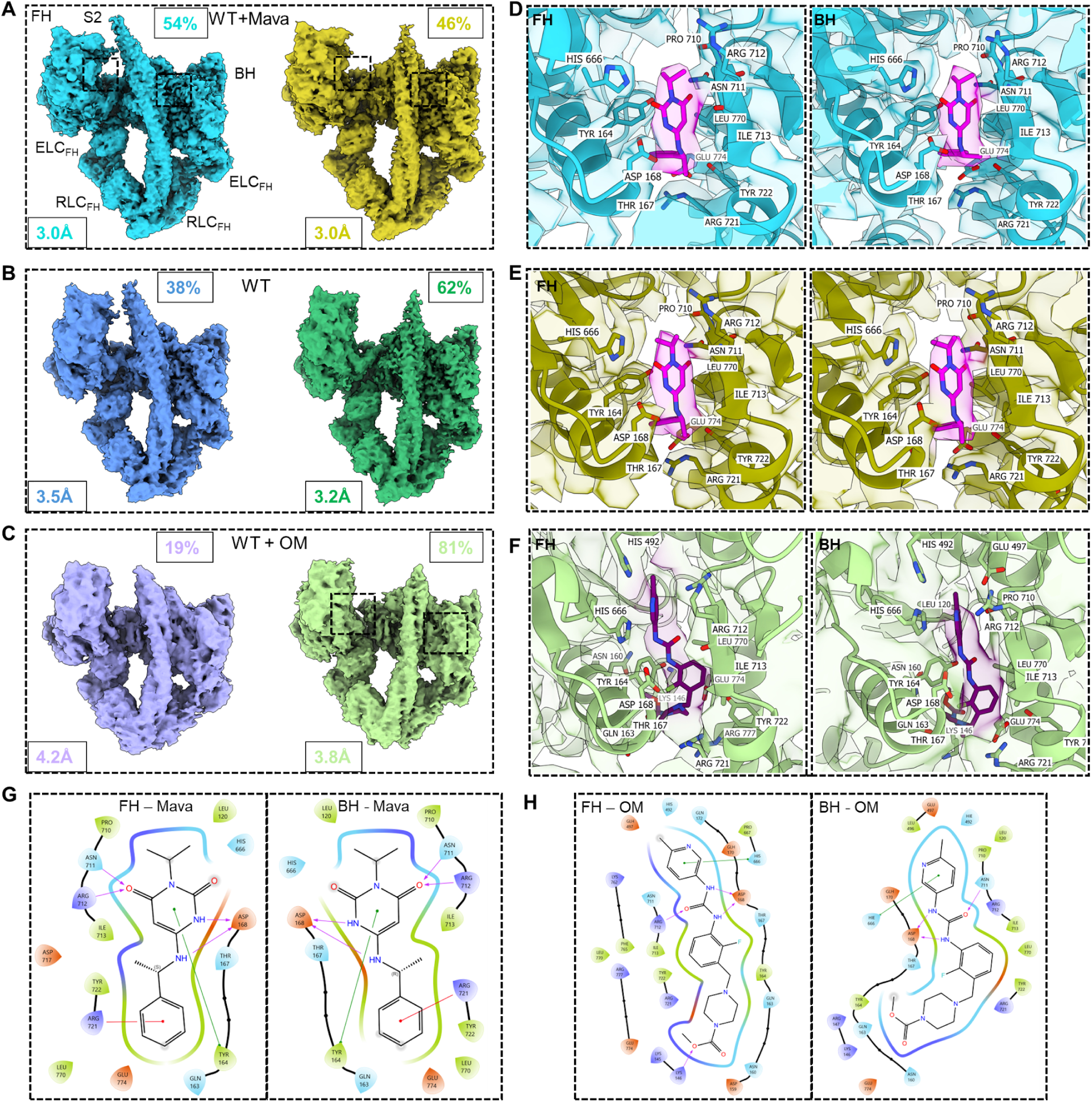
Cryo-EM density maps of the human β-cardiac myosin IHM under three experimental conditions. (A–C) M2β HMM with mava (A), without drug (B), and with OM (C). Each condition reveals two major IHM conformational states: a docked state (left images, shades of blue), where the S2 region bends and interacts with the FH, and an undocked state (right images, shades of green), where S2 is displaced from the FH and stays straight. 3D classification analysis (percentages shown for docked and undocked structures, see Tables S2 and S3 with statistical analysis) quantify the relative populations of the two states, showing that Mava favoring the docked and OM the undocked conformation. FSC global resolution is shown at the bottom left of each map. (D–F) Density maps fitted with atomic models focusing on the drug-binding sites and surrounding residues within the blocked and free myosin heads. (D, E) Mava (Magenta) in the docked and undocked IHM conformation binds to a conserved allosteric pocket, stabilizing (left, FH; right, BH) interdomain interactions. (F) OM binding site in the undocked IHM conformation showing OM (Purple) binding in both heads in the same allosteric binding pocket as Mava. (G, H) schematic ligand interaction diagrams of Mava (G) and OM (H) binding sites.

Mutations in β-cardiac myosin are one of the most common causes of inherited cardiomyopathies, with ∼30-40% linked to hypertrophic cardiomyopathy (HCM) and 3-4% to dilated cardiomyopathy (DCM)(*17, 18*). HCM-linked mutations often destabilize the IHM, increasing the population of myosin heads in the DRX state, thereby enhancing actin interactions and leading to hypercontractility(*19–22*). In contrast, DCM-associated mutations may stabilize the IHM, reducing actin interactions and resulting in hypocontractility(*22–24*). HCM and DCM mutations are thought to alter critical structural interactions in the myosin head-head, head-S2, and loop regions, altering the balance between active (open) and inactive (closed) conformations(*22, 25*).

Recent pharmacological advancements in the discovery of small molecules that can treat cardiomyopathies by directly targeting β-cardiac myosin are very promising(*26*). The myosin inhibitor Mavacamten (Mava, an FDA approved drug for treatment of obstructive HCM patients(*27*)) and the activator Omecamtiv mecarbil (OM, in phase 3 clinical trials for the treatment of DCM(*28*)), have shown potential in modulating the activity of β-cardiac myosin in diseased patients(*28, 29*). OM appears to destabilize the sequestered, energy-conserving IHM state, thereby increasing the population of heads available for actin interaction and force generation(*30*), thus enhancing myosin’s contractile force. In contrast, Mava enhances stabilization of this same sequestered IHM conformation, effectively reducing the number of active myosin heads and limiting hypercontractile activity(*31*). Despite the opposite effects of these two molecules on force production and heart contractility, x-ray crystallographic studies of isolated (single) myosin heads (S1) show that both target the same allosteric binding pocket (*32*). However, there have been no comparable structural studies on the (double-headed) IHM—the functional form of myosin in living muscle, having numerous interactions absent from S1(*13*). Therefore the structural mechanisms through which these drugs modulate IHM conformation remain unclear. Studying the high-resolution structural basis for the opposing effects of mavacamten and omecamtiv mecarbil is essential for guiding future drug design.

To understand the structural basis of these small molecule modulators, we here used cryo-EM to study high-resolution structures of human β-cardiac myosin in the sequestered IHM conformation both in the presence and absence of Mava and OM. Strikingly, in all three cases (WT, WT+Mava and WT+OM), we found two distinct conformations of the S2 region of the tail: docked and undocked. While Mava increases the percentage of conformations where S2 is docked onto the free head, stabilizing the IHM state, OM promotes undocked conformations, shifting S2 away from the FH towards a less stable IHM state. These changes align with their therapeutic roles in modulating contractility. Cryo-EM analysis shows in addition that while the docked conformation is stabilized by head-head and loop2-S2 interactions, these interactions are highly reduced in the undocked conformations. Our study provides a structural framework for understanding how these therapeutic drugs modulate the β-cardiac myosin IHM conformation. These insights could aid in developing next-generation modulators that precisely tune the stability of the sequestered IHM state to restore physiological cardiac function in diseased states.

## Results

### High-resolution cryo-EM structures reveal two distinct S2 conformations in the IHM

We employed single-particle cryo-EM to investigate the structural dynamics of an expressed human β-cardiac myosin IHM construct in the apo state (absence of drug, denoted wild-type, or WT) and under the influence of Mavacamten and Omecamtiv Mecarbil. Our IHM construct (M2β HMM) comprised the human β-cardiac MHC, with two heads and 25 heptads of tail, mouse skeletal light chains (ELC and RLC), a leucine zipper, and a C-terminal GFP tag(*24*) (Methods). The double-headed construct allows head-head and head-tail interactions, which are essential for forming the IHM conformation. 2D classification of the cryo-EM datasets identified IHM particles with well-defined structural features suitable for 3D reconstruction (Fig. S1-S3). In both WT and WT+Mava the IHM adopted a compact, well-ordered structure, consistent with previous cryo-EM observations (PDB ID: 8ACT(*13*), 9GZ1(*33*)); however, the IHM from WT+OM adopted a unique, less stable, distorted IHM architecture which will be discussed in detail later.

Following initial 3D reconstruction of each structure, we performed 3D classification to determine whether different conformations might underlie the averaged structures that were obtained using the entire dataset in each case. Strikingly, two distinct conformations of the S2 region were found across all three conditions (Fig. 1A-C), with S2 either bent, and docked with the FH (left images), or straight, and undocked (right images). Overall global resolutions of the structures were 3.0 to 4.2 Å (Fig. S4, Table S1), according to the gold-standard Fourier shell correlation 0.143 criterion. High-resolution density maps of WT and WT+Mava constructs revealed detailed structural features, including clear densities for ADP.Pi within the nucleotide-binding pockets of both the blocked and free heads in both docked and undocked structures (Fig. S5) and, in the case of the WT+Mava (3 Å resolution), clear densities for Mava are resolved in both heads of each structure (Fig. 1D, E). The WT+OM maps had lower resolutions (docked: 4.2 Å; undocked: 3.8 Å) but a modest OM density was still observed in the (more common) undocked state (Fig. 1F). With both OM and mava, the drugs occupied the same allosteric pocket, located between the converter region and the upper 50 kDa domain, as found in X-ray crystallographic studies of S1(*32*). This site is predominantly hydrophobic, engaging broadly conserved residues with non-polar side chains. Within 4 Å of each ligand, key interacting residues included N711, R712, I713, L770, E774, R721, Y722, T167, D168, Y164, and H666 (Fig. 1D-F).

Among these, the side chains of D168 (negatively charged) and N711 (polar), along with the backbone amide nitrogen of R712, formed direct hydrogen bonds with both ligands (Fig. 1G, H). Mava is a more compact molecule, primarily interacting with residues lining the central hydrophobic pocket, stabilizing binding through a combination of polar contacts and hydrophobic interactions. In addition to these Mava-interacting residues, the more elongated OM extended to reach H492 and E497 on the relay helix at one end and K146 at the other end, consistent with interactions observed in single-head bovine S1 structures(*32*). In the case of WT+Mava, the S2 density was resolved in sufficient detail to accommodate most side chains and visualize its coiled-coil architecture, revealing the heptad repeat organization, with hydrophobic residues forming the buried core and polar/charged residues exposed to solvent(*34*) (Fig. S6).

### Small molecule modulators alter the relative populations between the two S2 conformations

The relative population of the HMM between closed (IHM) and open structures, based on automatic blob-picking analysis, was dramatically impacted by the presence or absence and type of drug (Table S2). In the apo condition, ∼24 % of the molecules were classified as closed IHM, while ∼76 % corresponded to open, non-interacting heads. Mava shifted this equilibrium towards the closed state, increasing its population to ∼63 %, supporting the mechanism by which Mava reduces cardiac contractility through stabilization of the sequestered, energy-conserving IHM state(*31*). In contrast, OM induced a remarkable destabilization of the IHM conformation, with only 10 % of particles adopting the closed form and ∼90% as open, single-headed structures (similar to the FH of the IHM, which we interpret as an open, non-interacting heads structure (DRX state)).

Among the closed (IHM) particles, the S2 region exhibited the two distinct conformations (docked and undocked; Fig. 1) whose relative populations were also modulated by drug binding (Table S3; Fig. 1). In the apo condition, the IHM ensemble was composed of approximately 38% of particles in the docked and 62% in the undocked conformation (Fig. 1B, Table S3). In the presence of Mava, the conformational equilibrium shifted towards the docked state, increasing to 54% of the population, indicating enhanced stability of the IHM. This stabilization likely arises from strengthened interactions not only at the S2-FH interface, but also the S2-BH and BH-FH interfaces, which are more extensive in the docked conformation (discussed in detail later). By contrast, in presence of OM, among the small 10% IHM population, ∼80% of particles adopted the undocked conformation, indicating a strong shift towards a destabilized IHM state (Table S3).

### Structural basis of the two IHM conformations and their modulation by Mava and OM

The docked and undocked conformations of S2 are stabilized by two sets of related but differing interactions. In the docked state, S2 interacts at two crucial sites: first, with loop 2 of the blocked head, and second, where its bent C-terminal region docks on to the free head loop 2. In the second conformation, S2 is undocked from the FH loop 2 and adopts a straight, unbent orientation, while still maintaining interaction with loop 2 of the BH (Figs. 2, 3A, C).

**Fig 2.**
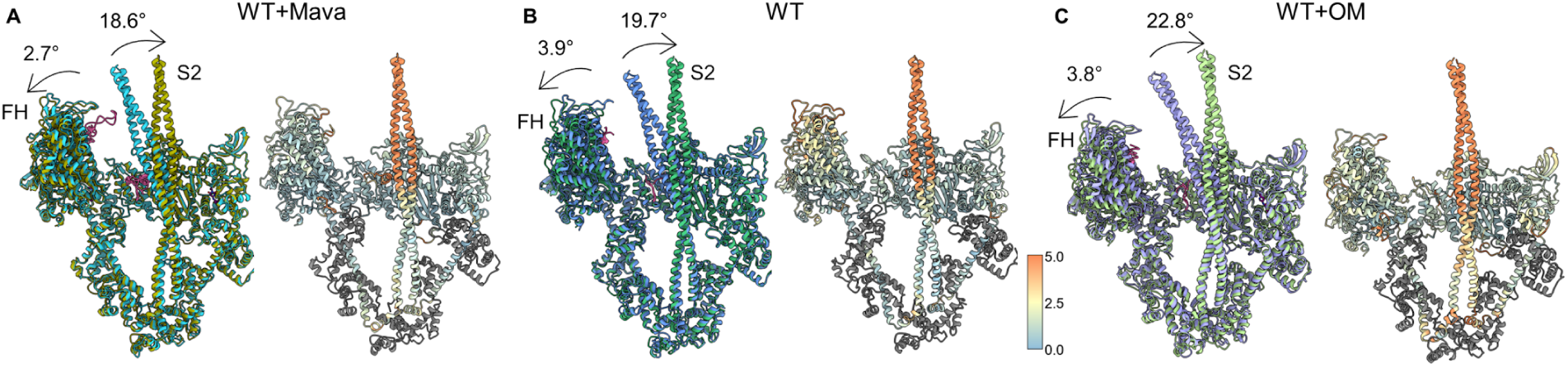
Displacement of S2 and the free head between bent and straight S2 conformations for the three experimental conditions. (A–C) Structural alignment of docked and undocked IHM conformations from WT+Mava (A), WT (B), and WT+OM (C) datasets. Within each panel, the docked (blue shades) and undocked (green shades) conformations are superimposed by aligning the BH to highlight the angular displacements of the FH (arrows to the left) and S2 (arrows to the right). Arrows indicate the direction and magnitude of structural shifts as S2 moves from docked to undocked position. These shifts reflect destabilization of the BH-FH interface as S2 undocks from the FH in each of the different conditions. While Mava favors minimal movement compared to WT, OM promotes a larger structural rearrangement (Table S4). Adjacent to each alignment, the corresponding undocked conformation is displayed and color-coded by per-residue RMSD relative to the docked state, mapped from low (blue) to high (orange) deviation, further showcasing the structural deviation in the S2 and FH regions. The bottom (yellow) region of the RMSD map, below the more strongly deviated orange region, represents the fulcrum of the bending movement of S2.

**Fig 3.**
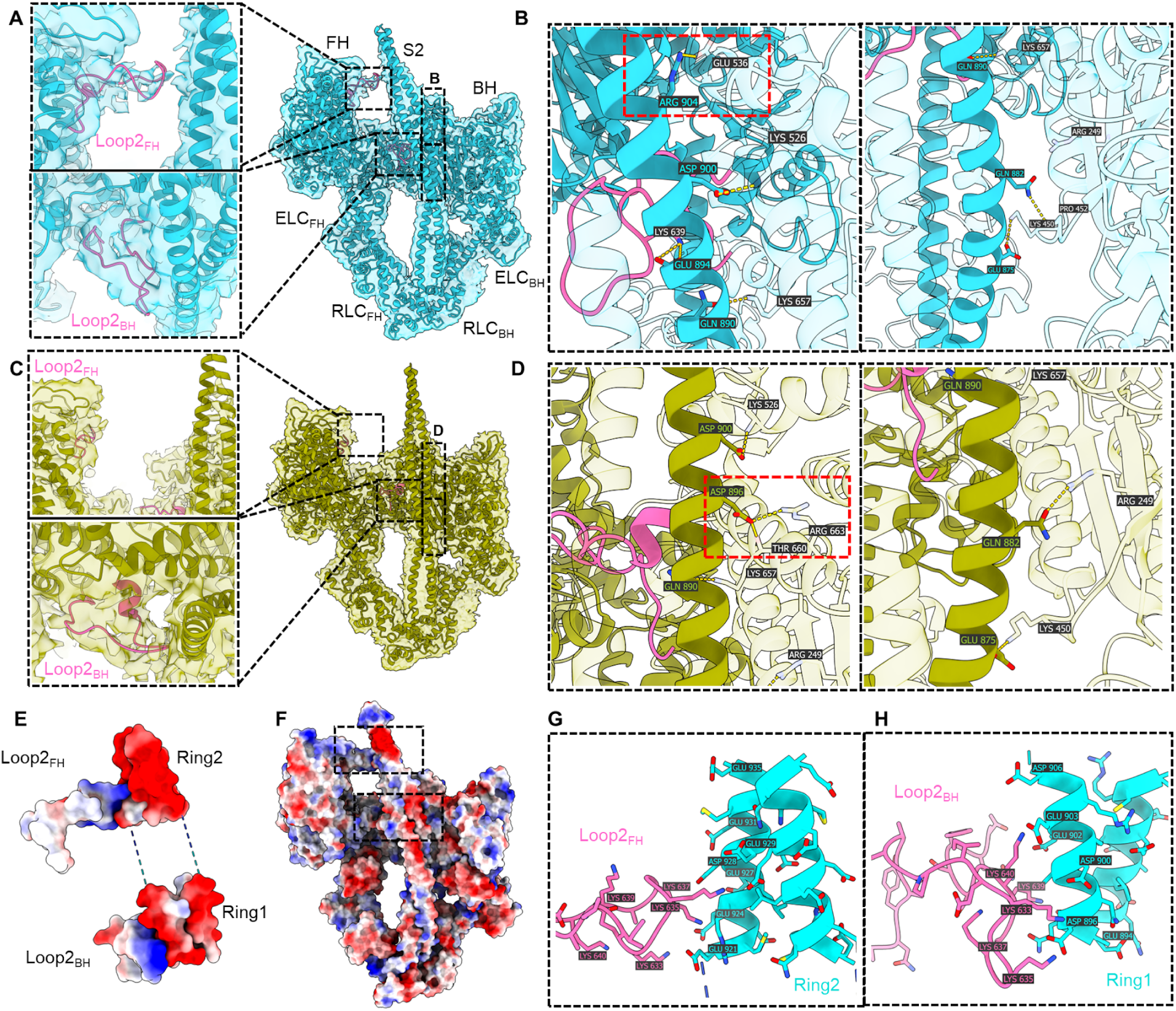
Cryo-EM density maps and atomic models of the WT+Mava (A,B) docked and (C,D) undocked IHM conformations. (A,C) Zoom in views show the interaction between Loop 2 (pink) of the BH and FH with different (lower and upper, respectively) regions of S2. (B,D) Zoomed-in panels of S2 – mesa interfaces showing S2 (from FH) interacting with positively charged residues from BH mesa, forming a triad of electrostatic interactions that strengthens the IHM configuration. Red-dotted rectangles show unique electrostatic interactions R904-E536 and D896-R663, possible only in the docked and undocked conformations respectively due to different positions of S2. (E, F) Electrostatic surface representation of WT+Mava docked IHM structure, highlighting charge complementarity regions between the motor domains and coiled-coil S2 region. Key interaction zones between Loop 2 of BH and Ring 1 of S2, and between Loop 2 of FH and Ring 2 of S2 (lower and upper rectangles respectively), showing well-aligned positive (blue) and negative (red) surface potentials. (G) Detail of FH Loop 2 – S2 Ring 2 interface, showing putative interacting charged residues, positive on Loop 2 and negative in Ring 2 of S2. (H) BH Loop 2 – S2 Ring 1 interface showing electrostatic interactions between basic Loop 2 residues and acidic residues of Ring 1.

In addition to movement of S2 between the docked and undocked positions, we have also observed a movement of the FH away from the BH, when S2 is undocked, causing a weaker BH-FH interface for the S2 undocked conformations. These synchronized movements highlight the dynamic nature of the S2 and FH regions and their pivotal role in mediating the BH-FH interactions within the IHM. Similar distinct S2 and FH conformations were observed in all three conditions. To better understand the structural transitions between the two S2 conformations, we measured the angular displacements of both the S2 and the FH regions relative to their positions in the docked state (Fig. 2). These measurements revealed distinct rotational movements of S2 that correlate with the strength of the BH-FH interface and help explain the structural basis for IHM stabilization or destabilization. Within each condition (WT, WT+Mava, WT+OM), the angular shift of S2 between docked and undocked conformations was: 19.7° in WT, 18.6° in WT+Mava, and 22.8° in WT+OM (Fig. 2). This displacement reflects the motion of the distal portion of S2 as it moves away from its contact site near FH loop 2 of the docked state, effectively breaking one of the key IHM interfaces (Fig. 3A, C and Movie S1). Simultaneously, the whole FH region undergoes a rotation between docked and undocked conformations of 3.9° in WT, 2.7° in WT+Mava, and 3.8° in WT+OM. This FH displacement represents a reorientation of the FH relative to the BH, weakening the head-head contacts crucial for maintaining the IHM state.

We also analyzed the positions of docked S2 in WT compared with WT+Mava and WT+OM (Table S4, col. 2). We made similar comparisons for undocked S2 and for the position of the FH in the docked and undocked S2 structures (Table S4). The S2 and FH positions were very similar (only 2-3° different) between WT and WT+Mava, suggesting that Mava maintains the IHM architecture close to that of the apo WT state. In contrast, the angular differences were much larger (∼10–12°) between WT and WT+OM states for both S2 and FH positions (Table S4), suggesting that OM substantially remodels the IHM architecture through structural destabilization of the S2–FH, FH-BH and S2-BH interfaces (discussed in detail later). These angular measurements demonstrate that both S2 and the FH undergo synchronized reorientation with drug treatment. While Mava stabilizes native interactions and maintains overall IHM architecture, OM causes substantial disruption of IHM structure, consistent with its role in enhancing active, open (non-IHM) conformations and thereby contractility.

### Loop 2 – S2–Mesa electrostatic interactions drive the stability of the IHM across drug conditions

As seen above, our cryo-EM reconstructions of the β-cardiac myosin construct under different conditions (WT, WT+Mava, and WT+OM) reveal a potential role of the S2 domain in regulating the stability of the IHM. To further understand the structural basis of this regulation, we examined the high-resolution maps from the WT+Mava dataset, which provided the most structurally resolved IHM (3.0 Å resolution) among the three conditions (Fig. 3). This allowed us to clearly visualize the two major conformational states, each characterized by distinct head-head and head-S2 electrostatic interactions.

The S2 region of the β-cardiac myosin IHM is formed by intertwining of the α-helical regions extending from the C-terminus of each head. We observed a striking charge-based complementarity at the S2–head interfaces, which help stabilize the IHM conformation (Fig. 3). On the side of the S2 facing Loop 2 of both the BH and FH, it is largely the S2 helix from the BH, which contains negatively charged residues (E and D) in the form of ring1(*35*) (residues E894 to D906) and ring2 (residues E921 to E935), that engages in electrostatic interactions with positively charged lysines (K633, K635, K637, K639, K640) in Loop 2 from both heads (Fig. 3A, C, E-H, Movie S1). On the opposite face of the coiled coil, the S2 helix from the FH, which has the same acidic residues (E and D), forms complementary interactions with basic residues (K and R) from the mesa(*21*) region of the BH (Fig. 3B, D). Some of the most notable contacts between S2_FH_ and the mesa region include salt bridges (E875–K450 and D900–K526) and polar hydrogen bonds (Q882–R249 and Q890–K657). In addition, depending on the S2 position, R904 can form a salt bridge with E536 in the docked state, and D896 forms a salt bridge with R663 in the undocked state. These two state-specific salt bridges are exclusive to those states and cannot form in the alternate conformation (Fig. 3B and 3D).

Comparison of the docked and undocked conformations in WT+Mava revealed that these interactions are substantially altered between the two states. In the S2-docked conformation, ring 1 of the S2_BH_ coiled coil interacts strongly with Loop 2 of the BH, while ring 2 of the S2_BH_ coiled coil bent towards the FH to interact with its Loop 2, forming a bridge that supports the FH positioning (Fig. 3E-H). This two-way interaction stabilizes the FH in this position, further strengthening the BH-FH interface and overall folded-back IHM architecture. These interactions are strikingly enhanced in the presence of Mava (through increasing the S2-FH docked population), correlating with its known function as a stabilizer of the sequestered state.

In contrast, in the S2-undocked conformation, we observed that the interaction between Ring 2 and FH loop 2 is disrupted, leading to collapse of the Loop 2_FH_–S2–Loop 2_BH_ bridge, causing detachment of the coiled-coil S2 region from the FH and making S2 straight. On the other hand, the contact between Ring 1 and BH loop 2 remains intact in the undocked state. This loss of S2-mediated interaction with the FH (Ring 2 and FH loop 2) correlates with a destabilization of the BH–FH interface and an increase in the conformational flexibility of the FH. While both docked and undocked conformations are observed across all three conditions, the IHM population in presence of OM is rarely observed, and when it is present, it is highly dominated by this weakened, undocked S2 conformation, consistent with its role in promoting active DRX states.

These dual-sided electrostatic interactions place the S2 in a precise orientation, bridging both motor domains in the docked state and contributing significantly to the structural integrity of the IHM. Loss of these interactions in the undocked states leads to S2 disengagement from the FH and rearrangement of the head-head geometry, thereby destabilizing the sequestered state (Movie S1). These structural changes are supported by the previously mentioned angular shifts in both S2 and FH (Fig. 2), again emphasizing a cooperative electrostatic interaction network involving S2, Loop 2, and the BH mesa. While Mava strengthens this precise interaction network, stabilizing the IHM, OM has a major disruptive effect, favoring active, open-head, force-generating states.

### Conformational dynamics of the BH – FH interface in regulating IHM stability

To investigate how the opposite angular shifts of S2 and the FH (Fig. 2) affect head-head interactions, we analyzed the BH – FH interface in detail. This is the prime region of interaction keeping the IHM conformation intact. The angular displacements of S2 and the FH are accompanied by substantial changes in the buried surface area (BSA) at the BH–FH interface (Fig. 4B), which correspond directly to changes in the number of interatomic contacts and stability within the IHM.

**Fig 4.**
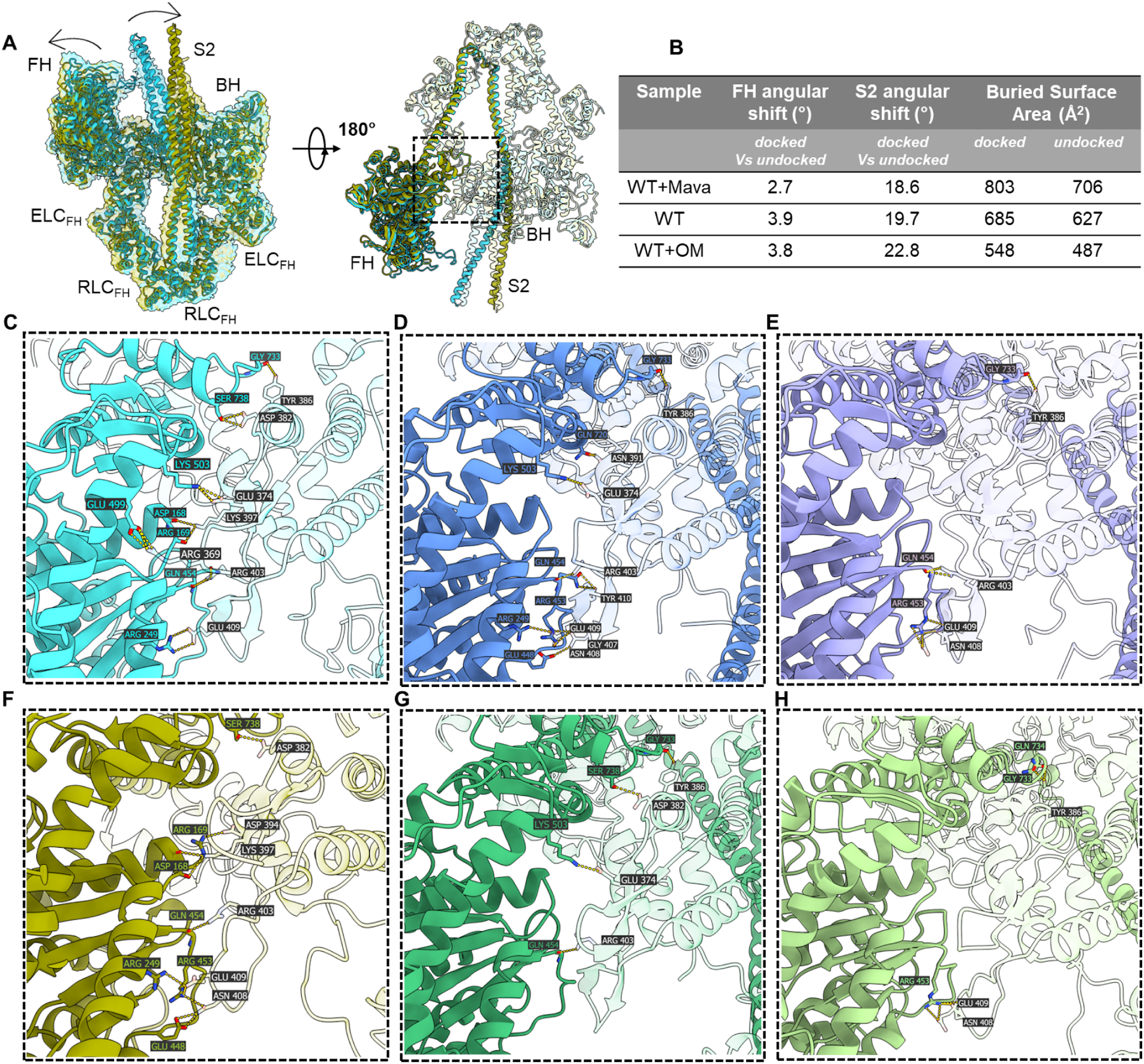
Comparative conformational dynamics and BH–FH interface interactions across WT, WT+Mava, and WT+OM. (A) Overlap of docked and undocked conformations of WT+Mava with their cryo-EM maps and models, showing angular displacements in the FH and S2 regions, and a 180° rotated view shows the BH–FH interface, indicated by the dotted box. (B) Quantitative comparison of FH and S2 angular shifts and buried surface area at the BH–FH interface. (C–E) BH–FH head-head interaction interfaces in docked conformations of WT+Mava, WT, and WT+OM, showing hydrogen-bonding interactions that stabilize the interface, with decreasing number of interactions from Mava to OM. (F–H) Corresponding views of undocked conformations reveal loss or rearrangement of these hydrogen bonds, with WT+OM showing the most distinct destabilization. To distinguish the BH and FH interacting residues, the BH is shown semi-transparent. Together, the comparisons illustrate how drug binding affects the BH–FH interface hydrogen bonding network and differentially modulates IHM stability across the variants.

The BH–FH BSA in the docked conformation is highest for WT+Mava (803 Å²), followed by WT (685 Å²), and lowest in WT+OM (548 Å²) (Fig. 4B), suggesting a gradient of interface stability across the three conditions. In the corresponding undocked states, the BH–FH interface is further compromised, with BSA reduced to 706 Å² in WT+Mava, 627 Å² in WT, and only 487 Å² in WT+OM. The reduction in buried surface area is especially striking in the WT+OM conditions (Fig. 4D,G vs. E,H), highlighting the potent allosteric destabilizing effect of S2 movement on the IHM interface. These reductions in BSA correlate well with the angular displacement of the FH away from the BH, supporting the interpretation that structural shifts in both the S2 and FH domains drive disassembly of the IHM.

To understand the underlying molecular interactions of these transitions, we analyzed specific inter-residue contacts at the BH–FH interface. In the docked conformation, a stable network of interactions, both electrostatic and polar hydrogen bonds, stabilizes the interface. The most prominent include BH-R369 – FH-E499, BH-E374 – FH-K503, BH-E409 – FH-R249 salt bridges (Fig. 4C). In contrast, the undocked conformation shows a pronounced loss of these interactions. At least two of the three key salt bridges observed in the docked state are disrupted, along with the reduction in BSA and an increase in FH conformational flexibility (Fig. 4B, F).

We next examined the molecular interactions at the BH–FH interface across WT (Fig. 4D, G) and WT+OM (Fig. 4E, H) conformations. In the WT docked state, the network of interactions ensures tight head-head association, supporting the compact and energy-conserving nature of this conformation. In the undocked conformations, particularly in the OM-bound conditions, we observed a progressive loss of these key salt bridges and contact residues, accompanied by a reduction in buried surface area to ∼ 548 Å² and 487 Å² respectively for docked and undocked states, indicating a weak head-head association in both docked and undocked states (Fig. 4E, H).

In summary, the transition from the docked to undocked IHM conformations is accompanied by the weakening or loss of key electrostatic interactions at the BH–FH interface. This loss, involving charged and polar residues across both heads, leads to electrostatic destabilization of the interface, resulting in increased conformational flexibility of the FH and weakening of the sequestered state. These effects are particularly pronounced in the WT+OM condition, consistent with the observed destabilization of the IHM state.

### Drug induced structural changes of the lever arm reveal allosteric control of IHM stability

In addition to changes in the S2 and FH domains, we observed a prominent shift in the BH and FH lever arms, which connect the motor domains to the coiled-coil S2 in the IHM. Given its structural role in mediating communication between the motor domain and the tail region, the lever arm serves as an important mechanical and regulatory element in cardiac myosin function. We measured and compared the angular shifts of the lever arms across all three conditions to assess how small-molecule modulators influence IHM architecture through this domain. While the angular shifts of S2 and the FH between docked and undocked conformations were observed across all three conditions, the lever arm shifts appear to be distinctly drug-specific.

To derive lever arm changes independent of motor domain movement, we first aligned all three docked conformations (WT, WT+Mava, and WT+OM) on the motor domain of the BH. Compared to WT, the BH lever arm in WT+Mava showed a subtle displacement of 1.9° (Table S5). In contrast, the BH lever arm of WT+OM showed a moderate 5.6° shift relative to WT, and 5.1° relative to WT+Mava, indicating that OM induces a conformational rearrangement in the BH lever arm even in the docked state (Fig. 5A, Movie S2,3). Similarly, when we aligned structures on the FH motor domains, we observed further large lever arm deviations. Comparing WT and WT+Mava, the FH lever arm showed a moderate 5.3° angular shift, whereas the FH lever arm in WT+OM docked conformation was displaced substantially by 13.1° relative to WT and 11.2° compared to WT+Mava, pointing to OM’s distinct effect on the lever arm geometry (Fig. 5B). A similar trend in lever arm displacement was observed across the undocked conformations in all three conditions (Fig. 5C, D, Table S5, Movie S2,3). Notably, in both Mava and OM-bound states, the FH consistently exhibited larger angular deviations than its partner BH, suggesting that the FH serves as the principal structural component driving both stabilization and destabilization of the IHM. This observation aligns with the angular deviations measured within the FH itself between docked and undocked conformations, underscoring its central role in mediating the dynamic equilibrium between the two structural states.

**Fig 5.**
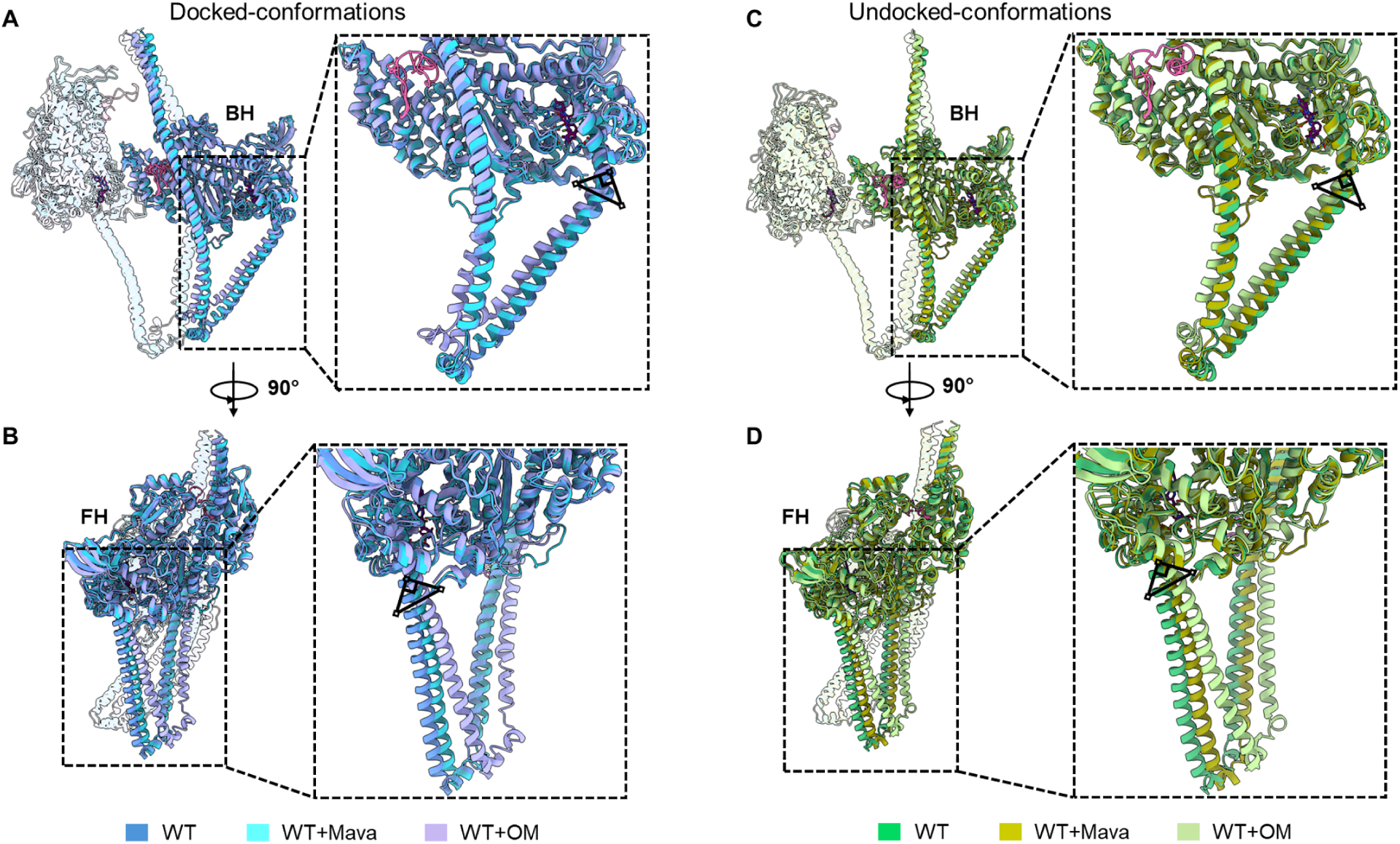
Lever arm angular deviations in docked and undocked IHM states upon drug binding. (A, B) Structural overlays of the three docked conformations (WT, WT+Mava, WT+OM) aligned on the blocked head (A) or the free head (B). Lever arm angular deviations for WT+Mava and WT+OM are measured relative to WT, which reveals drug-induced shifts in lever arm positioning (Table S5). (C-D) Structural overlays of the three undocked conformations (WT, WT+Mava, WT+OM) aligned on the blocked head (C) or free head (D). The small triangle depicts the fulcrum of lever arm deviation. Angular measurements demonstrate the differential effects of Mava and OM on lever arm orientation in both docked and undocked conformations.

In the presence of Mava, the lever arm undergoes a subtle to moderate angular rotation (∼2° for BH and 3-5° for FH) towards the upper 50 kDa region of the motor domain (Fig. 5B, D, Table S5). This facilitates the stabilization of inter-domain interactions between the U50 of the motor domain and the lever arm. In addition, this shift in the lever arm allows the formation of a key salt bridge between residues K146 and D778 in both docked and undocked conformations, and in both FH and BH (Fig. S7, Movie S2). The interaction is particularly significant in the FH, because this region is otherwise flexible and tends to displace away from the S2 segment. A subtle movement of the lever arm closer to U50 of the motor domain and formation of the K146–D778 salt bridge appears to restrain the lever arm, thereby minimizing FH flexibility and contributing to IHM stability, an observation which is also made recently by McMillan et al.(*33*) using a crosslinked HMM construct. In contrast, analysis of the apo WT structures reveals that these residues are unfavorably positioned to interact where the inter-residue distances of K146 and D778 are 3.6 Å and 5.8 Å in the docked and undocked conformations, respectively (Fig. S7E).

Upon structural alignment, we observed that angular shifts in the lever arm also induce significant and functionally relevant repositioning of the coiled-coil S2 domain across the BH mesa. Compared to WT, the lever arm displacement in WT+Mava resulted in only a minor adjustment, maintaining the relative orientation of S2 with respect to both motor domains (Movie S2). However, in the WT+OM condition, the pronounced lever arm remodeling caused the S2 to slide vertically across the BH mesa by nearly one helical turn (Fig. S8, Movie S4). This shift leads to a significant misalignment between Ring1 and BH loop2, as well as between Ring 2 and FH loop 2, disrupting the electrostatic network that normally stabilizes the S2–loop 2–mesa interface. The light chains, attached to the lever arm, also follow this large displacement, moving closer towards the motor domains and giving the overall appearance of a compressed or “squashed” IHM configuration. These major structural changes suggest that OM-induced lever arm reorientation not only alters the positioning of S2 but also disrupts the key interfaces (S2 – loop 2 – mesa) and inter-head geometry, highlighting the crucial role of the lever arm in allosteric modulation of IHM stability.

Together with our observations that angular displacements of S2 and FH correlate with weakening of the BH – FH interface and reduced buried surface area, these findings establish that drug induced remodeling of the lever arm is an essential part of the allosteric network governing IHM stability. While Mava strengthens a compact, energetically favorable IHM by minimizing lever arm flexibility, OM promotes a more distorted IHM conformation, facilitating transitions towards open active states. Thus the lever arm acts as a key structural component through which pharmacological drugs exert long-range effects on myosin regulation.

## Discussion

### Structural basis of cardiac myosin conformational flexibility

This study provides high-resolution structural insights into the conformational flexibility of the human cardiac β-myosin construct in its IHM state and reveals how small-molecule modulators influence its structural landscape. Using single-particle cryo-EM, we characterized distinct conformational ensembles of the IHM in WT, Mava-bound, and OM-bound states, and revealed key molecular mechanisms underlying drug-induced modulation of cardiac myosin.

One of the major observations across all three datasets is the dynamic behavior of the coiled-coil S2 region and its interactions with the FH. The observed IHM ensemble in each case is classified into two major conformations: a docked state with bent S2 interacting with the FH, and an undocked state with straight S2 displaced from the FH. While the docked conformations display an increased buried surface area and key stabilizing salt bridges at the FH-S2, BH – FH, and S2-loop2 interfaces, these interactions are strikingly reduced in the undocked state, making the IHM less stable. Computational studies of the IHM using molecular dynamics further supports our structural interpretations, where replacement of ADP.Pi by dADP.Pi (activator) weakens interface contacts and increases conformational heterogeneity in the interacting heads motif(*36*). The dual-sided electrostatic network between S2, Loop 2 and the BH mesa explained here precisely orients the myosin heads supporting the folded-back IHM and stabilizing head-head geometry. Consistent with this structural analysis, previous ab initio models used cryo-EM reconstruction of the tarantula thick filament to interpret loop 2 – S2 interactions in cardiac myosin and suggested that Loop 2 could form electrostatic contacts with negatively charged rings on the S2, stabilizing the heads and maintaining the folded IHM conformation(*25, 35*). A recent study further proposed that loop 2 may transiently interact with the S2 Ring 2 region to form a pre-IHM state, further emphasizing the importance of these interactions in IHM assembly and stability(*37*).

### Functional implications for cardiac contractility

These docked and undocked conformational states are closely linked to functional consequences, including the transition between SRX and DRX states(*38*) (where the closed, IHM structure unfolds to an open, non-interacting heads conformation which can interact with actin), influencing energy consumption and force generation in cardiac muscle contraction. Mava stabilizes and promotes the equilibrium towards the S2-docked IHM state, reducing the number of heads available for actin binding and lowering energy consumption(*31*). OM favors the S2-undocked state, shifting the equilibrium towards activation-ready DRX-like states, increasing head availability and contractile output(*30*) (Fig. 6A, B). Our data suggest that the FH – S2 – BH mesa interaction network is not only a structural feature, but a significant element for overall IHM stability. Loss of S2 docking on the FH correlates with increased FH flexibility, a weakened BH–FH interface, and overall destabilization of the IHM, particularly in the presence of OM (Fig. 4B). This supports the hypothesis that stabilization of FH-S2 contacts along with the S2-BH mesa and BH-FH interfaces are essential for the IHM docked preservation. This view is consistent with the unifying hypothesis where most HCM-associated mutations lead to a destabilization of the IHM state either directly or allosterically, resulting in more myosin heads being available for actin interaction(*19, 39*), and a comprehensive mutational analyses in which HCM variants at IHM interfaces show destabilization of SRX(*22, 40*).

**Fig 6.**
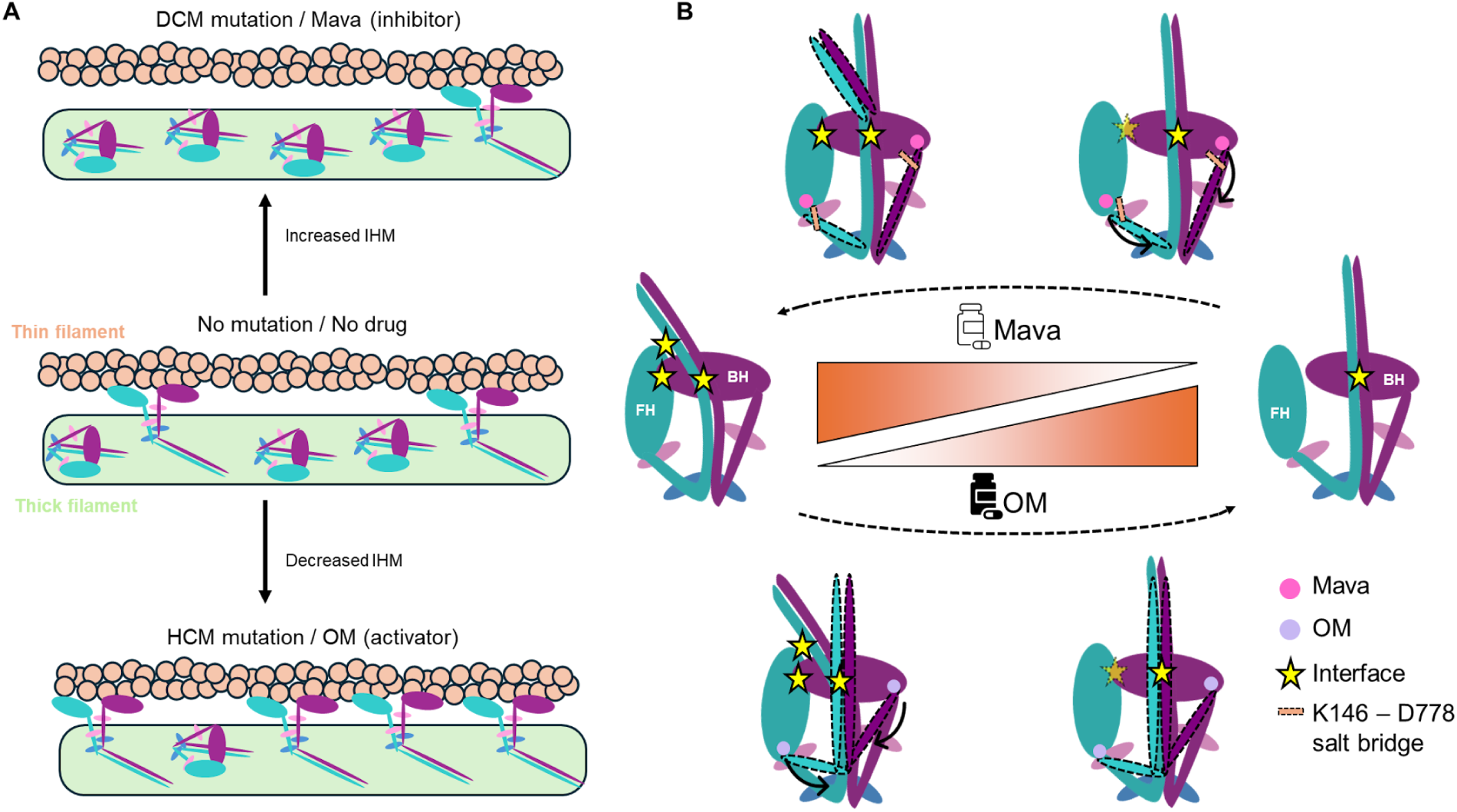
Proposed molecular mechanism for the allosteric control of IHM stability by Mava and OM(A). Schematic overview of thick and thin filament interactions illustrating how WT, HCM, DCM, and drug treatment shift the equilibrium between the sequestered IHM and open myosin states. In the normal WT heart (middle), both IHM and open states coexist in an optimal equilibrium. HCM mutations and OM shift the balance toward open myosin states, increasing actin-binding capability. DCM mutations and Mava shift the equilibrium toward increased IHM population, favoring energy conserving states. **(B).** Structural interpretation of IHM stability based on cryo-EM analysis. The docked state is maintained by three major interfaces: S2–FH, BH–FH, and S2–BH (yellow stars). Dotted-line ellipses and solid arrows highlight the transitions of the lever arm and S2 between docked and undocked conformations. Mava promotes lever arm shifting toward the motor domain, restricting lever arm flexibility and strengthening electrostatic interactions across the three interfaces, thereby stabilizing the bent S2 and docked IHM. In contrast, OM induces major leaver arm reorientation, weakening these interfaces and promoting the straight S2 and undocked conformation. Thus, Mava and OM allosterically modulate IHM stability by oppositely shifting the conformational equilibrium between docked and undocked states.

### Drug induced lever arm and interface modulation

Drug binding altered both lever arm orientation and interface stability in distinct ways. Mava stabilizes and increases the docked conformation population by inducing subtle reorientation in the lever arms towards the motor domains, restraining the lever arm flexibility, and strengthening electrostatic interactions at the S2 – loop 2 – mesa and BH-FH interfaces (Figs. 4 and 5). This helps in achieving the folded IHM conformation, consistent with previous structural and kinetic studies showing that Mava favors the SRX state by stabilizing the closed autoinhibited folded-back conformation of myosin heads(*31, 41*). In contrast, OM induces a major lever arm angular displacement towards the motor domain, weakening these interfaces and promoting the undocked conformation population (Figs. 4 and 5). This helps to release both the heads into an activation-ready conformation (Fig. 6B). Our WT+OM IHM structures provide the first visualization of OM-induced rearrangements of lever arm and S2 within the native two-headed conformation, extending previous S1-based observations that OM binds to an allosteric site and stabilizes the lever arm in a primed position thus increasing actin binding(*42–44*). The large lever-arm deviations observed in WT+OM (∼10° - 12°; Table S4, 5) further induced a considerable shift in the coiled-coil S2 position across the mesa compared to WT structures (Table S4 and Movie S3,4). This suggests that the lever arm can mechanically transmit allosteric drug bound signals from the motor domain to the distant structural regions such as S2.

### Comparison with previously reported IHM structures

Our IHM structures (global resolutions: WT, docked-3.5 Å, undocked-3.3 Å; WT+Mava, 3.0 Å, docked and undocked; WT+OM, docked-4.2 Å, undocked-3.8 Å), demonstrate both similarities and differences with other cryo-EM structures of WT (PDB 8ACT)(*13*) and WT+Mava (PDB 9GZ1)(*33*) (global resolutions of 3.6 Å and 3.7 Å, respectively). While the overall architecture of the IHM is conserved, our structures reveal a substantial degree of conformational variability, particularly in the positioning of the S2 and its interaction with the FH, that has not previously been reported, providing valuable insights into the dynamic behavior of the IHM. Strikingly, the S2 region of 9GZ1 aligns well with our docked conformation of WT+Mava, whereas S2 from 8ACT closely resembles our undocked conformation of WT (Fig. S9). The alternate positions of S2 revealed in our structures were not reported in either case.

A recent cryo-EM study of the human cardiac thick filament reveals IHMs in two distinct conformations, where the interacting motor domains (BH and FH) are either perpendicular to the filament axis (CrH) or tilted with respect to the filament axis (CrT; (*6*)). Comparison of our docked and undocked structures with these thick filament IHMs highlights the significance of these conformational states in the context of the native thick filament. Our docked IHM conformations align well with the CrT configuration, whereas the undocked states resemble the CrH conformation (Fig. S10). This correlation suggests that the conformational flexibility we observe at the single-molecule level underlies the distinct functional capabilities of these two configurations of IHM in the cardiac thick filament (*6*), underpinning insights into the molecular mechanisms by which the therapeutic drugs Mava and OM may function in the cardiac thick filament.

### Implications on pathogenic genetic mutations

Mapping of pathological HCM and DCM associated mutations onto the IHM, both isolated and in the thick filament, shows significant clustering at IHM interaction sites, including the mesa - S2 and BH-FH interfaces, which are crucial for IHM stability(*19, 22, 25, 40*). Many of these HCM mutations weaken BH–FH or S2–mesa interactions(*21, 22*). Notably R249Q, R663H, E536D, E374V and D382Y disrupt key electrostatic interactions at these interfaces, which would shift the equilibrium toward undocked, DRX-like states, thereby increasing the proportion of heads available for actin interaction, leading to hypercontractility. This is consistent with the unifying hypothesis of hypercontractility where HCM mutations cause increased contractility due to increased ON-state myosins(*19*). Strikingly, the DCM-associated mutation E525K, which causes decreased contractility, does the opposite. Its location at the BH-S2 interface appears to strengthen BH-S2 interaction through attraction of K525 in the mesa to negatively charged residues of Ring 2 in S2(*45*), stabilizing the IHM and enhancing the SRX state(*23, 24*). Other DCM variants such as S532P and F764L, affect myosin function by predominately altering mechanochemical properties of the motor domain(*46*).

Our data support the hypothesis that modulation of IHM stability is a central mechanism for fine tuning cardiac contractility, and may contribute to disease phenotypes. For example, mutations that weaken S2-mesa, S2-FH and BH-FH interactions may shift the balance toward the DRX state, increasing the number of heads available for contraction, causing hypercontractility as seen in HCM (Fig. 6A). Stabilization of the IHM, as seen with Mava, may help restore IHM like states, reducing the availability of heads and reducing pathological force generation (Fig. 6B). In contrast, conditions such as DCM may be treated by activators like OM that destabilize the IHM and increase the number of available heads, enhancing overall force output. Thus, our findings help map a structure-function relationship from molecular insights to pathological phenotypes.

## Methods

### Cryo-EM grid preparation

The human β-cardiac myosin 25 Hep (25 heptads of the coiled-coil) HMM construct comprised residues 1–1016 of the heavy chain. This was attached to a GCN4 leucine zipper (sequence: MKQLEDKVEELLSKNYHLENEVARLKKLVGER), a short linker (GSGKL), and a C-terminal arrangement of EGFP, Avi-tag (GLNDIFEAQKIEWHE), and FLAG-tag (DYKDDDDK) as previously described by Duno-Miranda et al(*23*). Purified WT HMM protein was diluted to a final concentration of 1.5 µM in a buffer containing 10 mM MOPS (pH 7.4), 20 mM KCl, 2 mM MgCl₂, 1 mM EGTA and 0.5 mM ATP. For drug-bound samples, Mava or OM was incubated with protein for 15 min at 25 °C before vitrification. 3 µl aliquots were applied to UltrAufoil R 1.2/1.3, 300 Mesh gold grids glow-discharged for 30 s at 25 mA using a PELCO easiGlow (Ted Pella). Grids were blotted for 4 s at 25 °C and 95% relative humidity using a Vitrobot Mark IV, followed immediately by plunge-freezing in liquid ethane. Vitrified grids were then stored in liquid nitrogen until screening and data collection.

### Cryo-EM data collection

Grids were initially screened on a 200 keV Talos Arctica to assess ice thickness, particle distribution, and orientation. Selected high-quality grids were later imaged on a 300 keV Titan Krios (Thermo Fisher Scientific) equipped with a Gatan K3 direct electron detector and a Gatan GIF Quantum energy filter. Automated data collection in counting mode was performed in multiple sessions using SerialEM v4.0(*47*), with separate datasets collected for each experimental condition. A total of 10363, 7070 and 11894 movies were recorded for WT, WT+Mava and WT+OM respectively. Each movie was composed of 40 frames over a 4 s exposure, corresponding to a total electron dose of ∼51 e⁻ Å⁻². Images were acquired at a nominal magnification of 105,000x, with a super-resolution pixel size of 0.415 Å (physical pixel size 0.83 Å). Data were collected with a defocus range of −1.0 to −2.0 μm.

## Data processing

All datasets from each experimental condition (WT, WT+Mava, WT+OM) were processed independently in cryoSPARC v4.6(*48*) (Structura Biotechnology). Movies were motion-corrected for beam-induced motion correction using Patch Motion Correction, followed by contrast transfer function (CTF) estimation using Patch CTF Estimation. Micrographs with high-quality CTF fits were selected for further analysis. Particle picking was performed using an initial automated blob picking approach followed by 2D classification to separate single motor domain open HMM states and to obtain initial IHM templates. These representative IHM templates were then used to train a neural network based particle picker Topaz(*49*) for enhanced detection of IHM particles. The particles from both Blob and Topaz picking were combined, and duplicates were removed using a distance cutoff of 100 Å. The resulting combined particle set was subjected to multiple rounds of 2D classification to remove false positives and poorly aligned particles.

For each dataset (WT, WT+Mava, and WT+OM), an ab initio reconstruction was performed with multiple classes, which served as references for heterogeneous refinement in cryoSPARC. Particles corresponding to well-defined 3D conformations were selected for homogeneous and non-uniform refinement. The resulting 3D model and particle set were subjected to 3D classification using a mask covering the entire IHM. The classes where clear densities were confirmed were separated into two distinct classes of maps (docked vs undocked conformations) based on the S2 and FH positioning.

The final 3D refinement and postprocessing of the classes yielded high-quality reconstruction maps with global resolutions of 3.0 Å (WT+Mava-docked), 3.0 Å (WT+Mava-undocked), 3.5 Å (WT-docked), 3.3 Å (WT-undocked), 4.2 Å (WT+OM-docked), 3.8 Å (WT+OM-undocked). Final global resolutions were determined using the gold-standard Fourier shell correlation (FSC = 0.143) criterion, and map sharpening was performed using B-factors automatically estimated by cryoSPARC. Local resolution maps were also calculated in cryoSPARC to assess density variation across structural regions. The outline of the processing strategy is described in Fig. S1-S3.

### Model building and refinement

Atomic models were built to interpret the cryo-EM density maps obtained for each experimental condition. An initial homology model of the human β-cardiac myosin heavy chain and mouse skeletal light chains was generated using the WT sequestered IHM structure (PDB ID: 8ACT(*13*)) as a template. The model was first rigid-body fitted into the docked and undocked cryo-EM maps using UCSF ChimeraX(*50*). The coiled-coil S2 segment was modeled using the crystal structure of the human β-myosin S2 fragment (PDB ID: 2FXM(*35*)). The entire IHM structure was then flexibly fitted into the density map using Namdinator(*51*), which allows for molecular dynamics based flexible fitting against the experimental map. Subsequent iterative real-space refinement was performed in PHENIX(*52*) with secondary structure, rotamer, and Ramachandran restraints applied. Manual adjustments, including loops and side chains, were performed in Coot(*53*) to improve local fitting. For drug-bound conditions, ligand geometry restraints were generated using eLBOW (PHENIX suite). Ligand coordinates for Mava and OM were placed into their respective binding pockets, refined using real-space refinement in PHENIX, and visually inspected to ensure optimal fit to the map density. Model validation was performed using MolProbity(*54*) analysis, geometry measurements, and model-to-map correlation provided by PHENIX.

## Acknowledgments

We thank C. Ouch, K. Song and C. Xu of the cryo-EM Facility at the UMass Chan Medical School for help and training in cryo-EM imaging; K. H. Lee and G. Hendricks of the Electron Microscopy Facility for conventional EM training; This work was supported by the National Institutes of Health, grant numbers HL164560, AR081941, HL163585, and HL150953.

## Author contributions

A.K.S. designed and conducted experiments, prepared cryo-EM grids, processed and analyzed cryo-EM data, and carried out reconstruction, atomic fitting, refinement, structure analysis, database deposition and wrote the first draft. J.G, C.M.Y contributed to myosin sample preparation and biochemical characterization. A.K.S., R.C and R.P. interpreted the data and wrote the manuscript. All authors discussed the results and approved the final version of the manuscript. Funding acquisition, project administration and supervision: C.M.Y, R.C. and R.P.

## Data availability

The cryo-EM density maps and corresponding atomic models have been deposited into EMDB and PDB under accession codes EMD-73268, 9YOP (WT docked), EMD-73367, 9YRG (WT undocked), EMD-73288, 9YP9 (WT+Mava docked), EMD-73362, 9YR7 (WT+Mava undocked), EMD-73283, 9YP4 (WT+OM docked), and EMD-73368, 9YRH (WT+OM undocked). PDB data used to build the models were 8ACT and 2FXM. In addition, the following cryo-EM map EMD-29722 and PDB models were used for comparisons: 8ACT, 9GZ1 and 8G4l

## Supplementary Materials

**Fig. S1.**
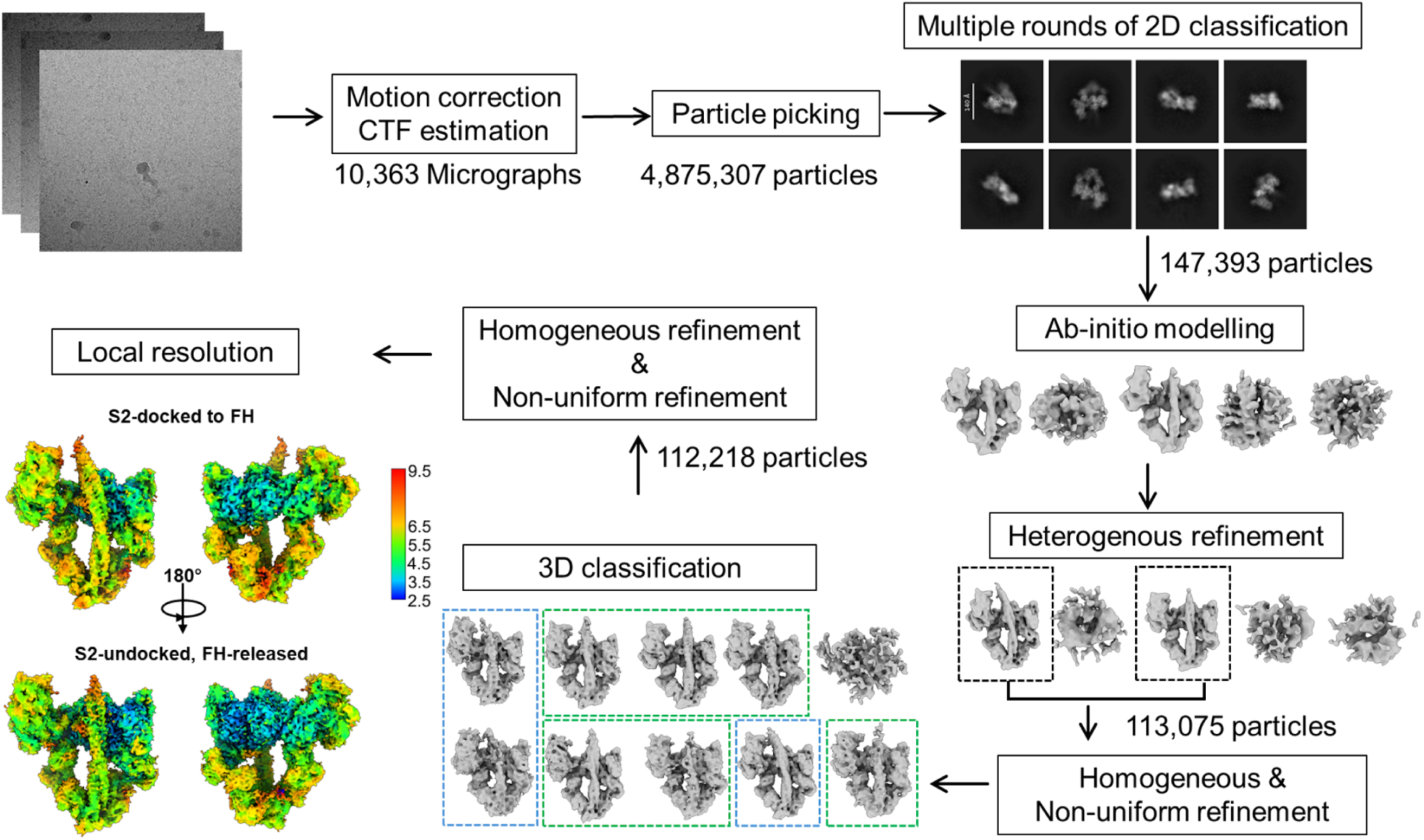
Cryo-EM data processing workflow of WT. Schematic overview of the image processing pipeline for WT, including motion correction, CTF estimation, blob and template-based particle picking, multiple rounds of 2D classification to separate single motor domains as well as junk particles from IHM particles, ab initio reconstruction and heterogeneous refinement for particle curation, and 3D classification leading to the final docked and undocked IHM maps.

**Fig. S2.**
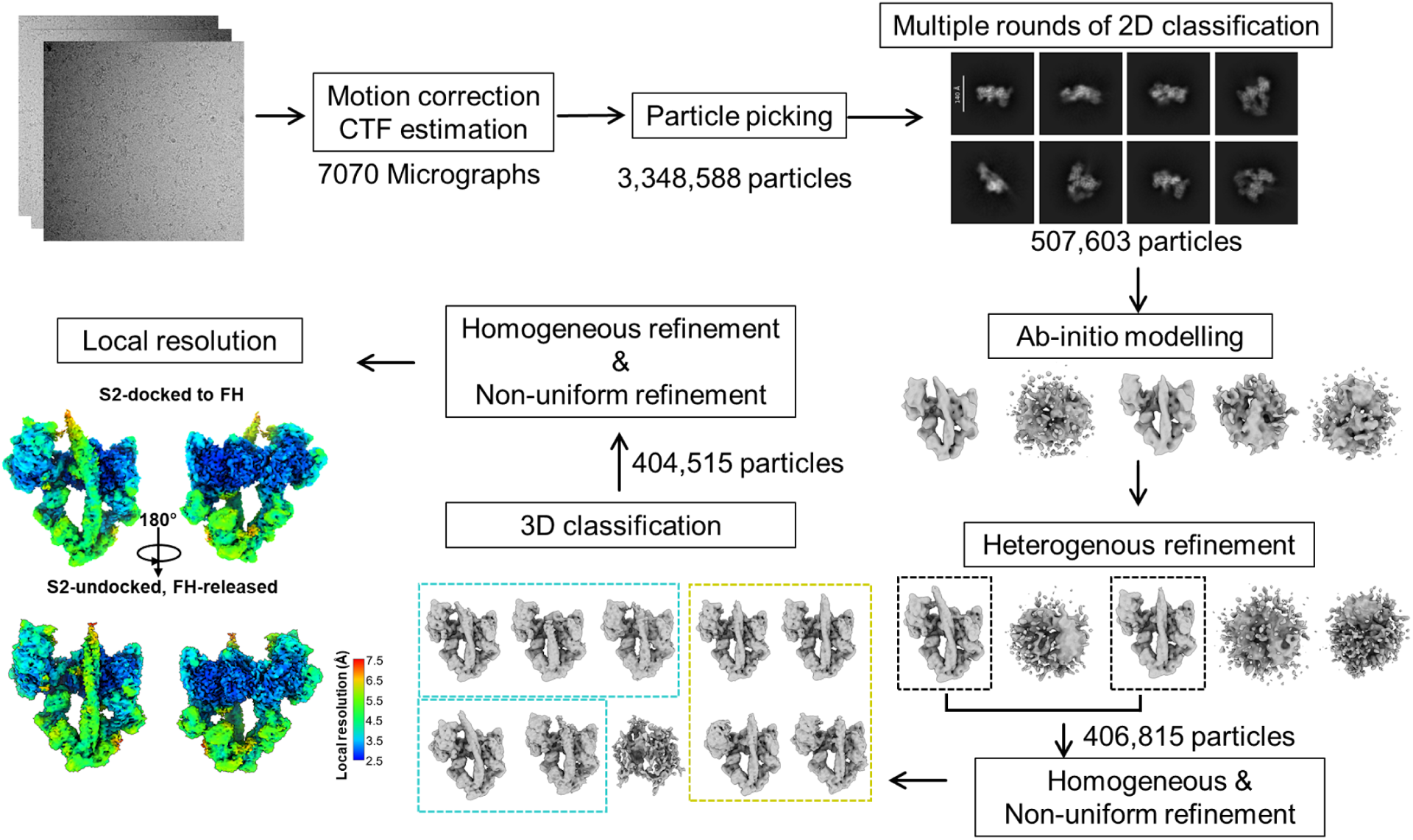
Cryo-EM data processing workflow of WT+Mava. Schematic overview of the image processing pipeline for WT+Mava, including motion correction, CTF estimation, blob and template-based particle picking, multiple rounds of 2D classification to separate single motor domains as well as junk particles from IHM particles, ab initio reconstruction and heterogeneous refinement for particle curation, and 3D classification leading to the final docked and undocked IHM maps.

**Fig. S3.**
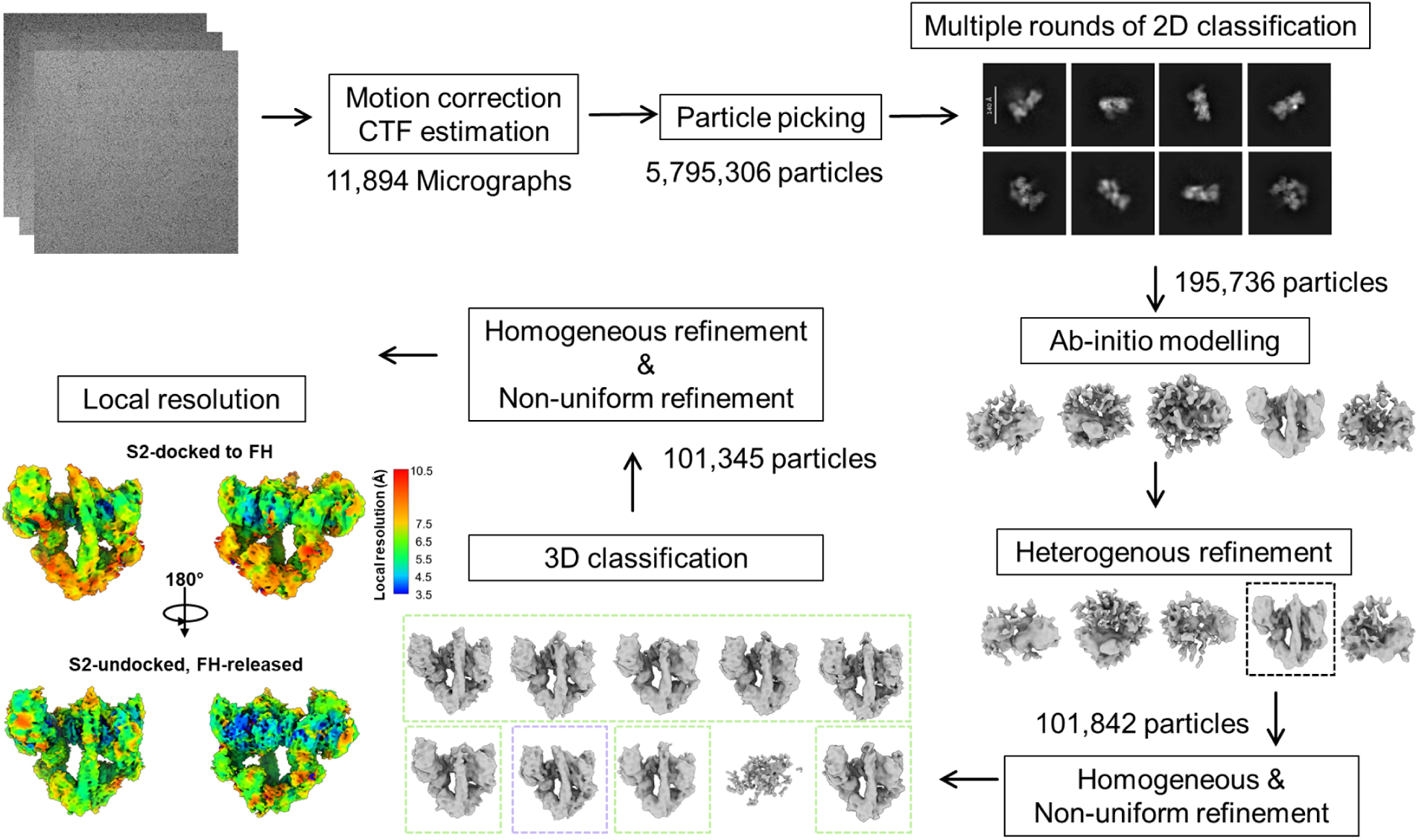
Cryo-EM data processing workflow of WT+OM. Schematic overview of the image processing pipeline for WT+OM, including motion correction, CTF estimation, blob and template-based particle picking, multiple rounds of 2D classification to separate single motor domains as well as junk particles from IHM particles, ab initio reconstruction and heterogeneous refinement for particle curation, and 3D classification leading to the final docked and undocked IHM maps.

**Fig. S4.**
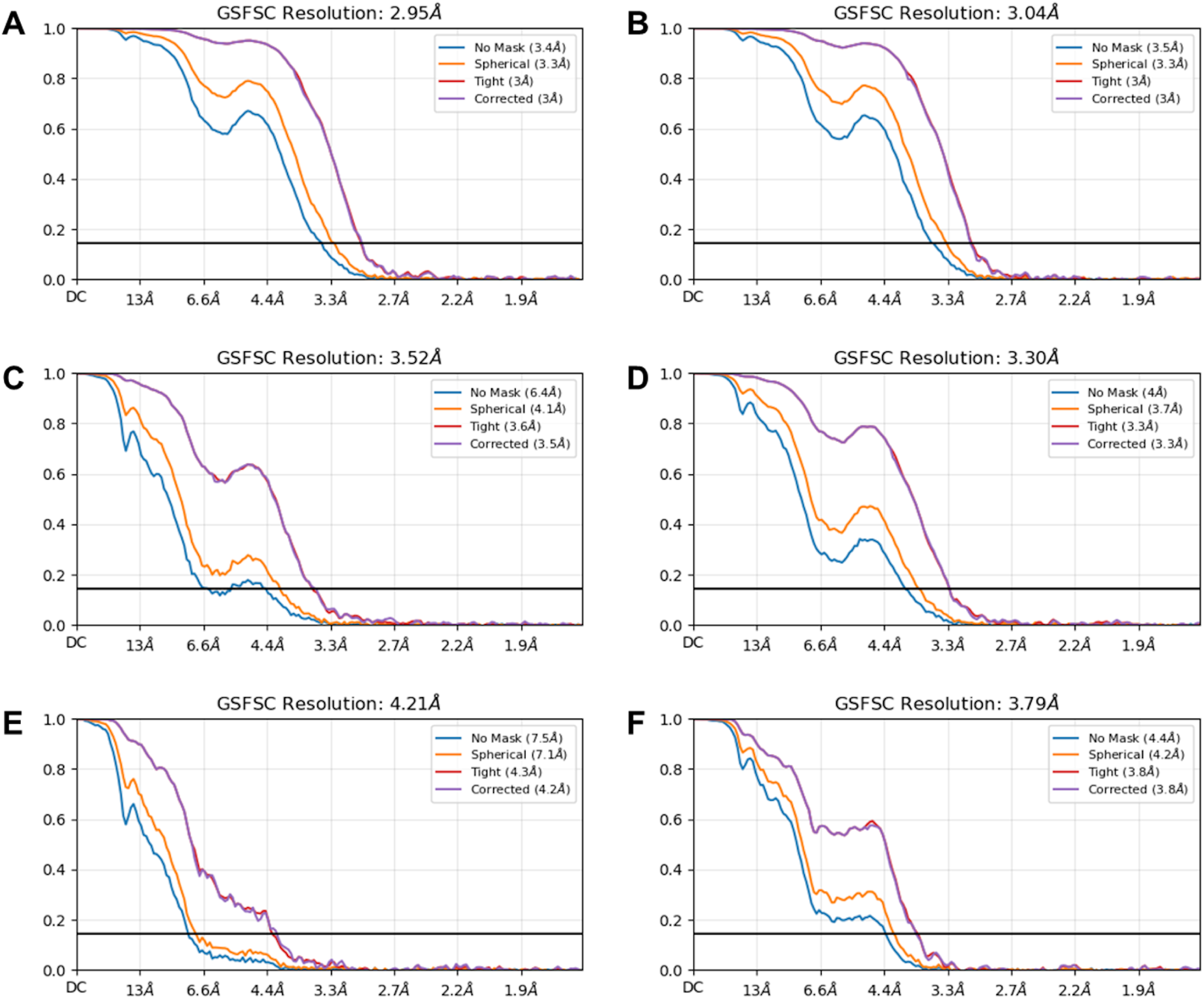
Gold-standard Fourier shell correlation (FSC) plots for the final 3D reconstruction maps. FSC plots are shown for WT+Mava (A, B), WT (C, D), and WT+OM (E, F) datasets, separated into docked (A, C, E) and undocked (B, D, F) conformations. The FSC = 0.143 threshold is indicated by the black line.

**Fig. S5.**
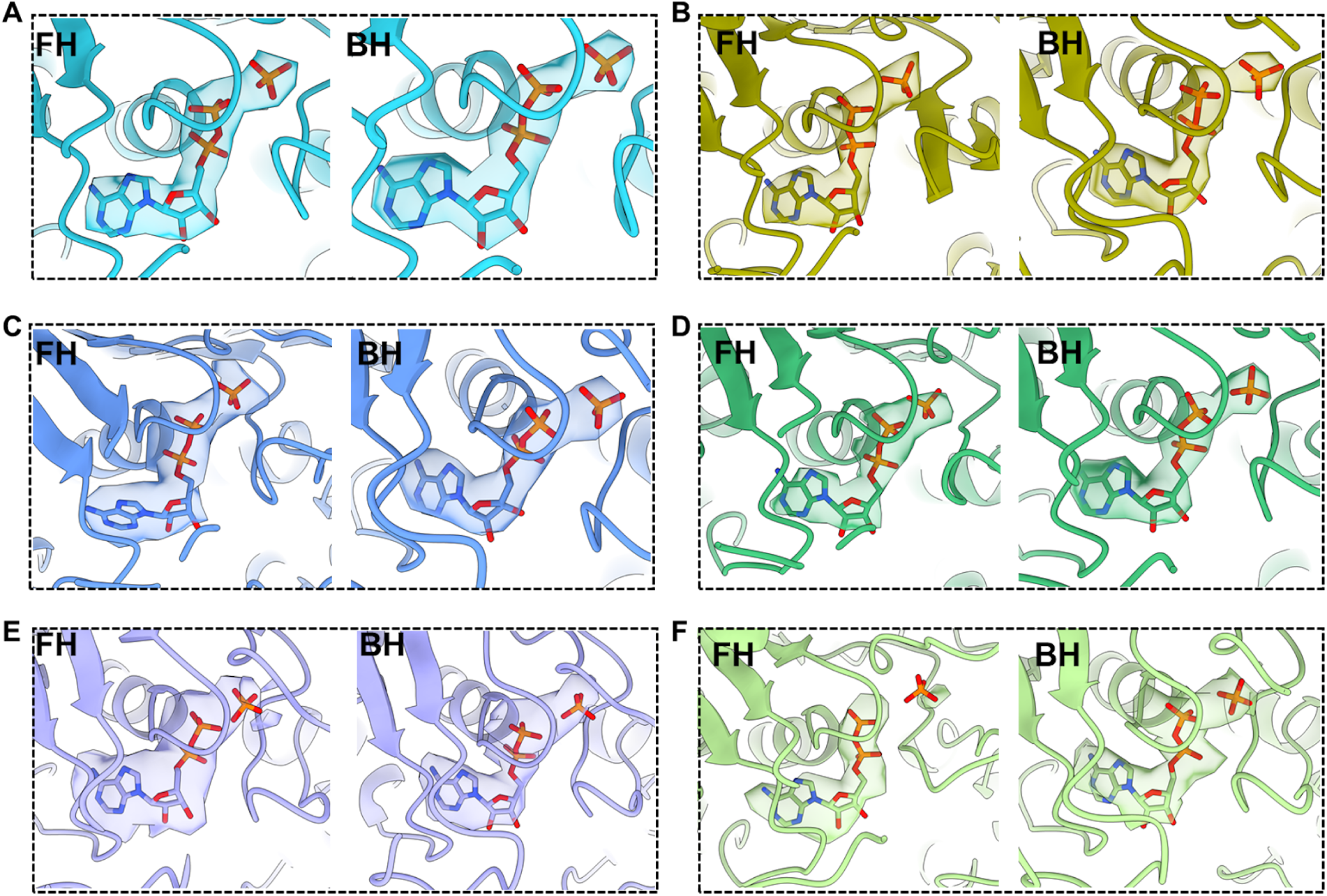
Fitting of ADP·Pi into cryo-EM densities of WT, WT+Mava, and WT+OM IHM conformations. (A, B) Segmented cryo-EM density maps of WT+Mava in the docked (A) and undocked (B) conformations, showing placement of ADP·Pi within the nucleotide binding pockets of both the BH and FH. (C, D) Equivalent fitting for WT in the docked (C) and undocked (D) conformations. (E, F) Segmented densities of WT+OM in the docked (E) and undocked (F) conformations with ADP·Pi fitted into the corresponding BH and FH sites. In every case, with both BH and FH, ATP is seen to be in the hydrolyzed state, with Pi separated from ADP.

**Fig. S6.**
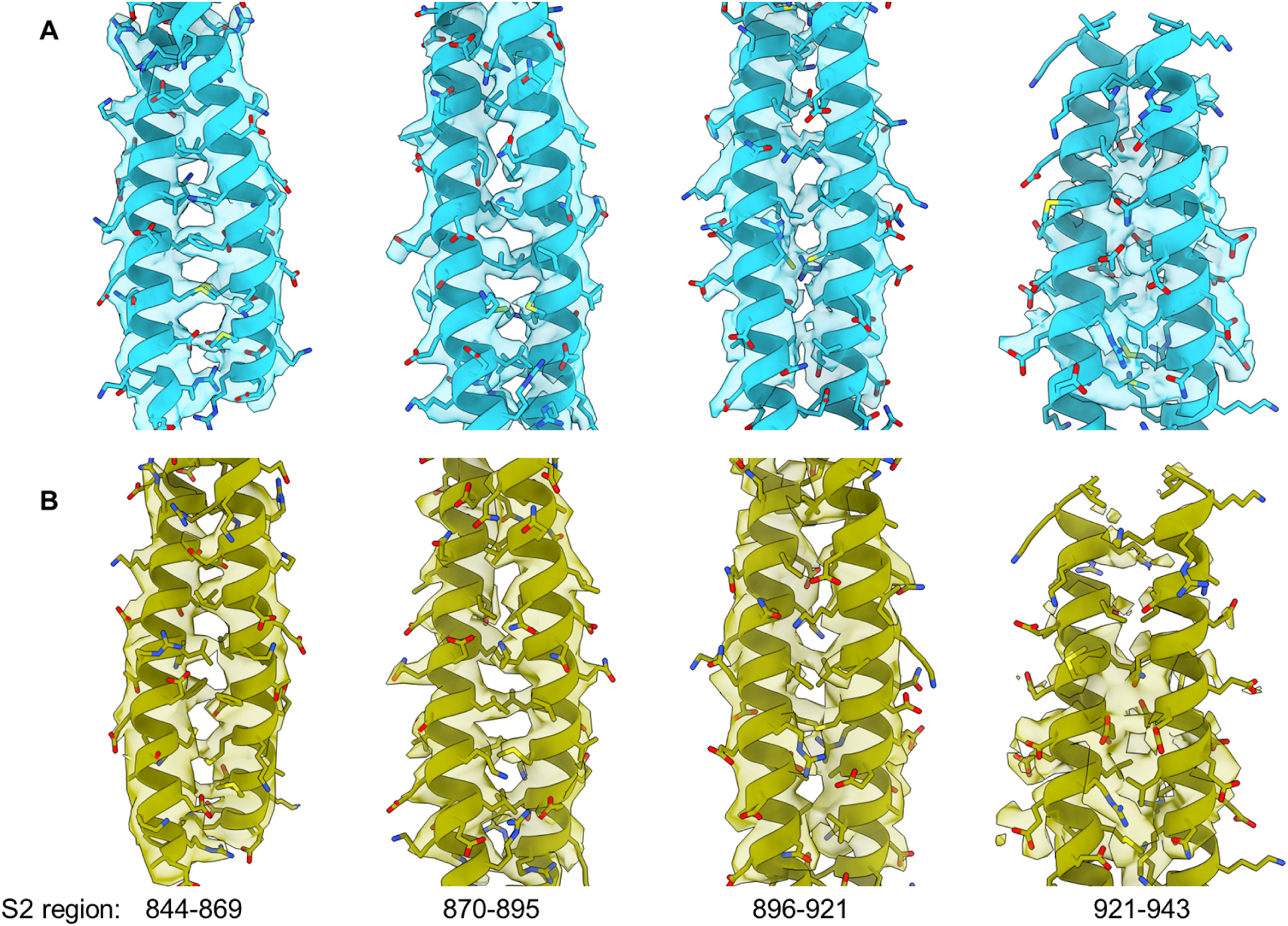
Coiled-coil arrangement of the flexible S2 region resolved in both the docked (A) and undocked (B) cryo-EM maps of WT+Mava. The segmented cryo-EM density of the S2 coiled-coil region of residues 844–943 fitted with the atomic model was divided into four panels. This clear density, accommodating the majority of side chains, shows the standard coiled-coil arrangement where the hydrophobic residues align along the heptad to form the buried core, while the charged and polar residues are exposed to solvent.

**Fig. S7.**
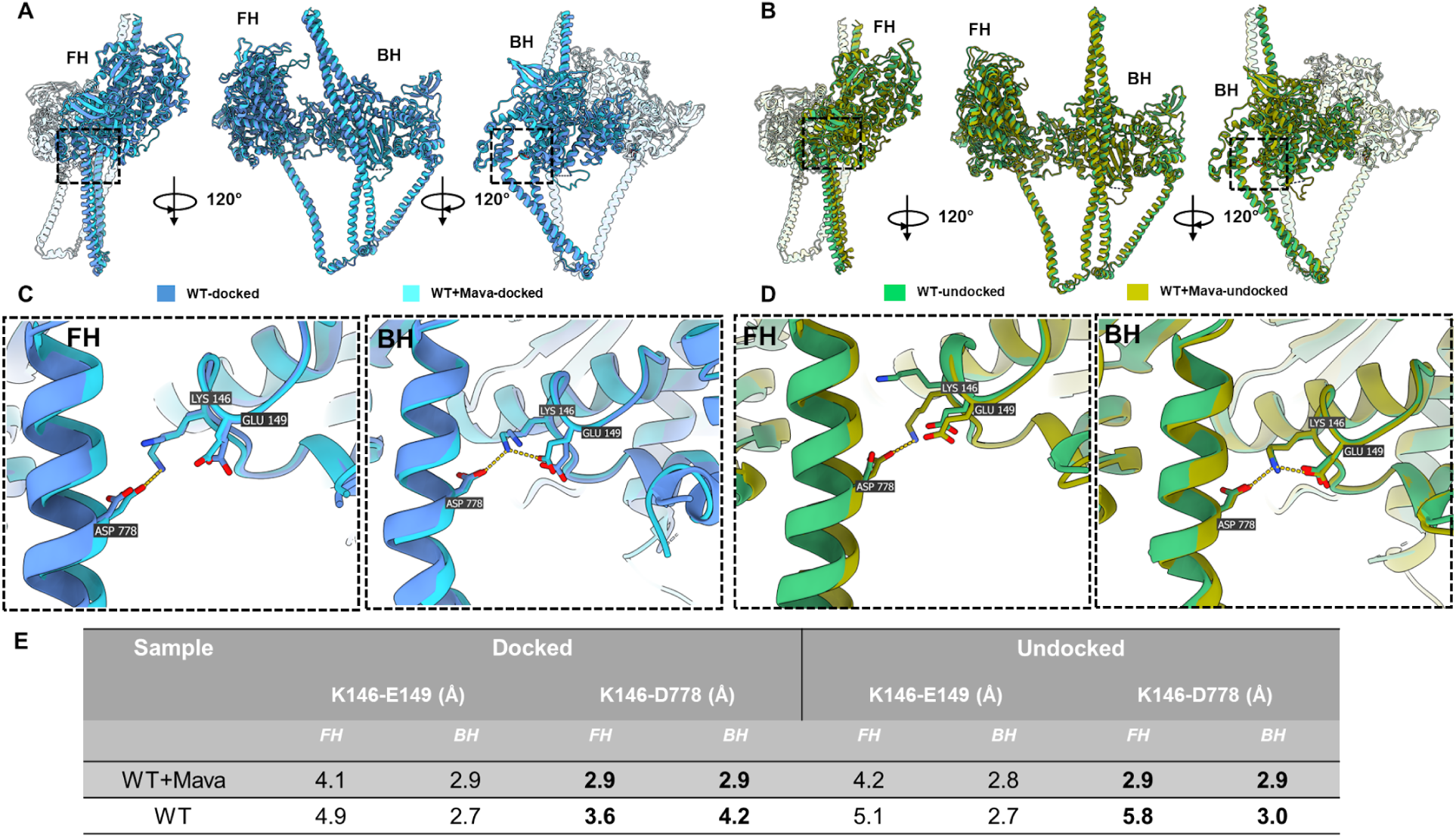
Mava restrains lever arm movement and reduces FH flexibility, contributing to IHM stability. (A, B) Comparison of WT and WT+Mava IHM conformations aligned on the blocked head in the docked (A) and undocked (B) states. (C, D) Close-up views of the K146–D778 salt bridge region in both heads for docked (C) and undocked (D) conformations. (E) Overall, in the BH, K146 lies between E149 and D778 at a distance of ∼3.0 Å, where it can potentially interact with either residue. However, in the FH, K146 does not interact with E149 but instead forms a salt bridge with D778 in the lever arm, particularly in WT+Mava.

**Fig. S8.**
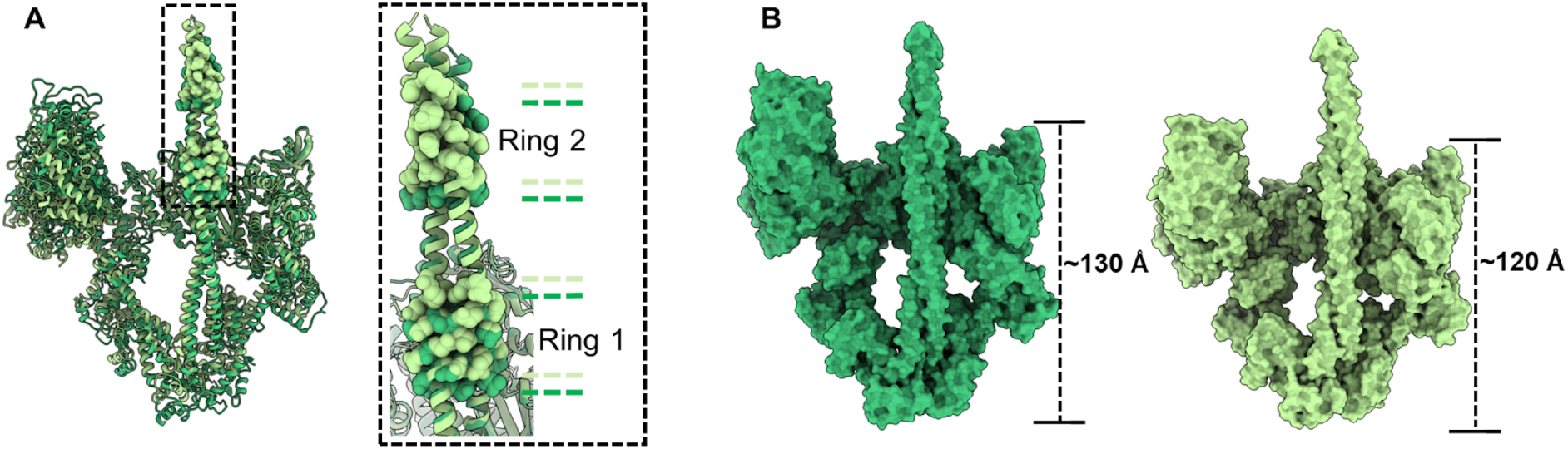
OM-induced displacement of the S2 region and lever arm remodeling in the undocked IHM. (A) Structural comparison of WT undocked and WT+OM undocked (the most populated state) conformations aligned on the BH, highlighting OM-induced repositioning of the S2 domain. The displacement of S2 is visualized by labeling Ring 1 and Ring 2 regions as spheres, showing their vertical shift across the BH mesa upon OM binding. (B) Surface representation comparing the WT undocked and WT+OM undocked conformations. In WT undocked, the BH extends nearly 130 Å along the S2 axis, whereas in WT+OM undocked, the length decreases to ∼120 Å due to the pronounced lever arm angular shift and compression of the IHM.

**Fig. S9.**
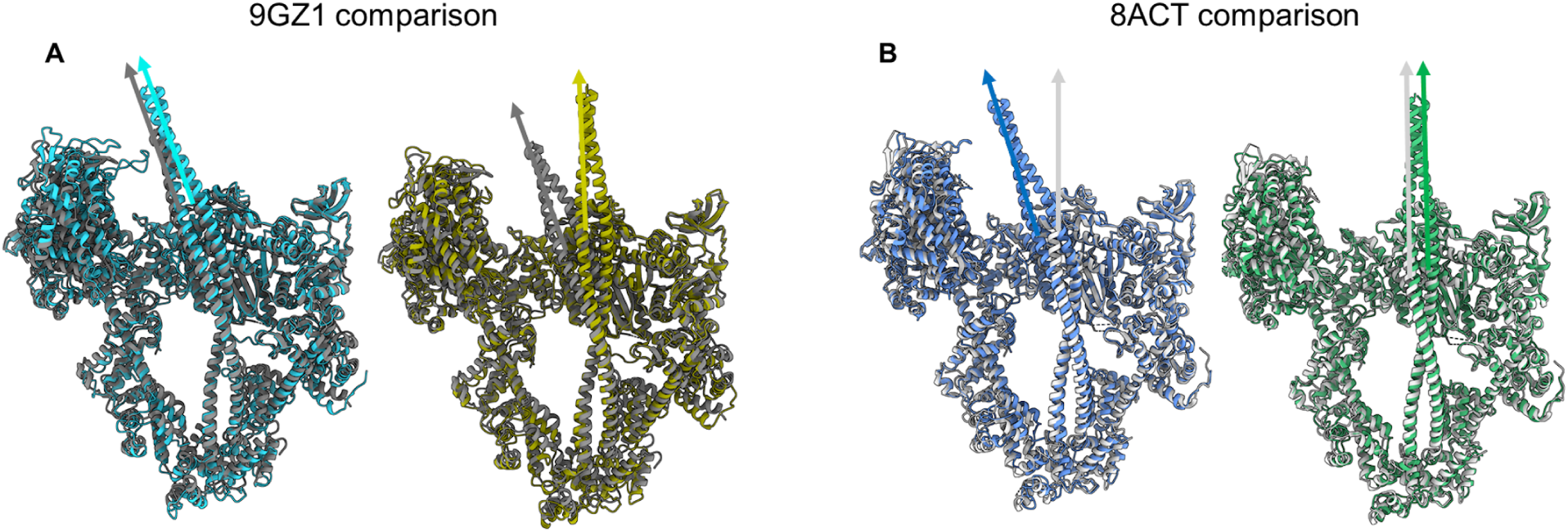
Comparison of recently published IHM structures (grey) with our cryo-EM reconstructions (colors). (A) Comparison of PDB 9GZ1 (WT+Mava) with our WT+Mava docked and undocked conformations shows that 9GZ1 most closely resembles the docked state, where the S2 is bent towards the FH, which was the more populated in our study. (B) Comparison of PDB 8ACT (WT) with our WT docked and undocked conformations shows that 8ACT most closely corresponds to our more populated WT undocked state, where the S2 stays straight.

**Fig. S10.**
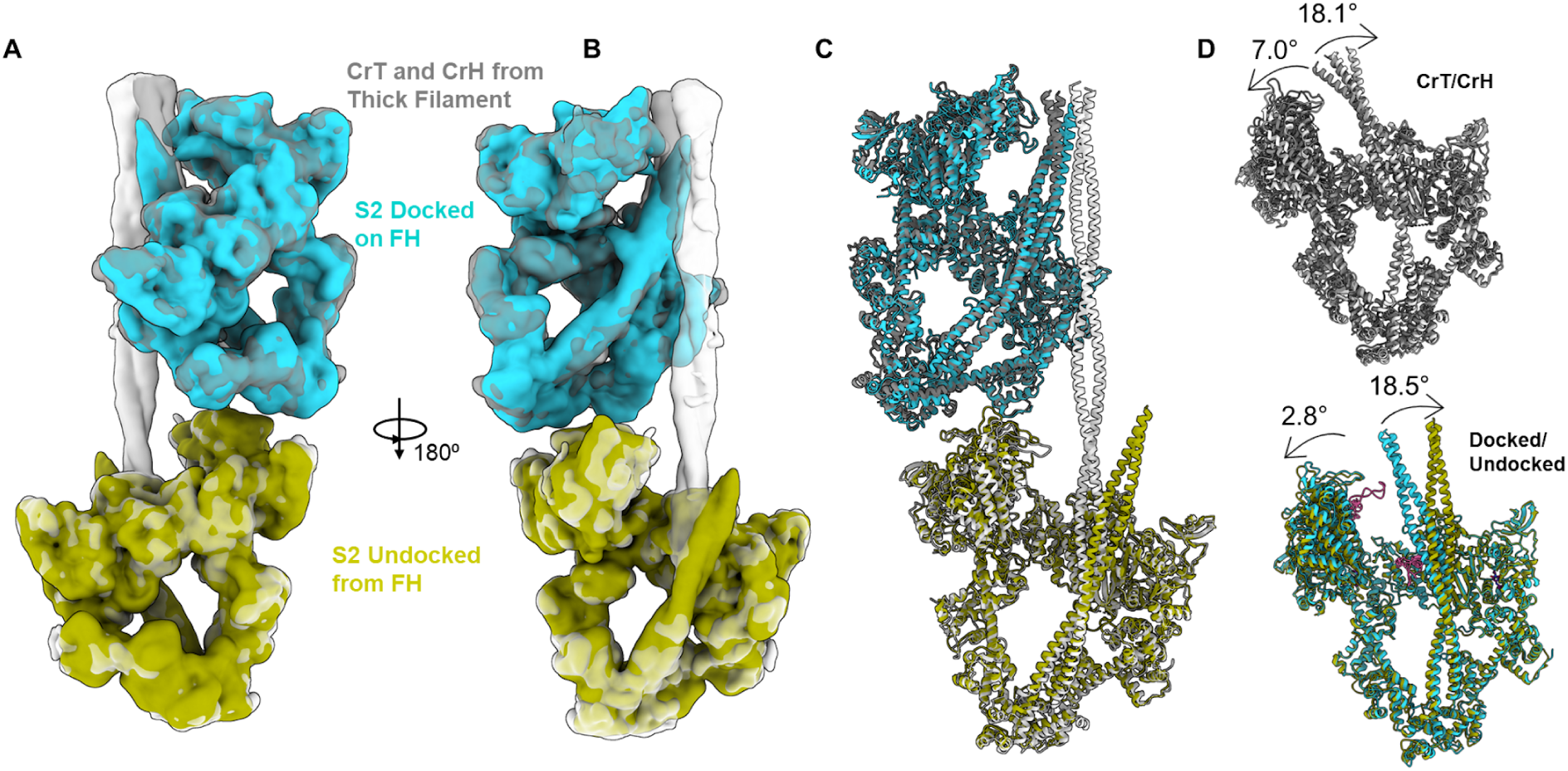
Comparison of WT+Mava IHM conformations with thick filament structures. (A, B) Fitting of WT+Mava docked and undocked density maps (blue, green) into the CrT and CrH segmented maps (grey) from the cardiac thick filament reconstruction, also in the presence of mava (Dutta et al., 2023). Our single particle WT+Mava reconstructions were low-pass filtered to 6 Å to match the resolution of the thick filament map. (C, D) Structural comparison of WT+Mava docked and undocked atomic models with the CrT and CrH thick filament IHMs from the atomic model PDB 8G4L, respectively. Alignments were performed on the blocked head, and angular displacements of the FH and S2 domain were measured (see Fig. 2A). The comparison reveals similarities in FH positioning and S2 orientation between the single particle IHM conformations and the thick filament IHM states in different crowns.

**Table S1.**
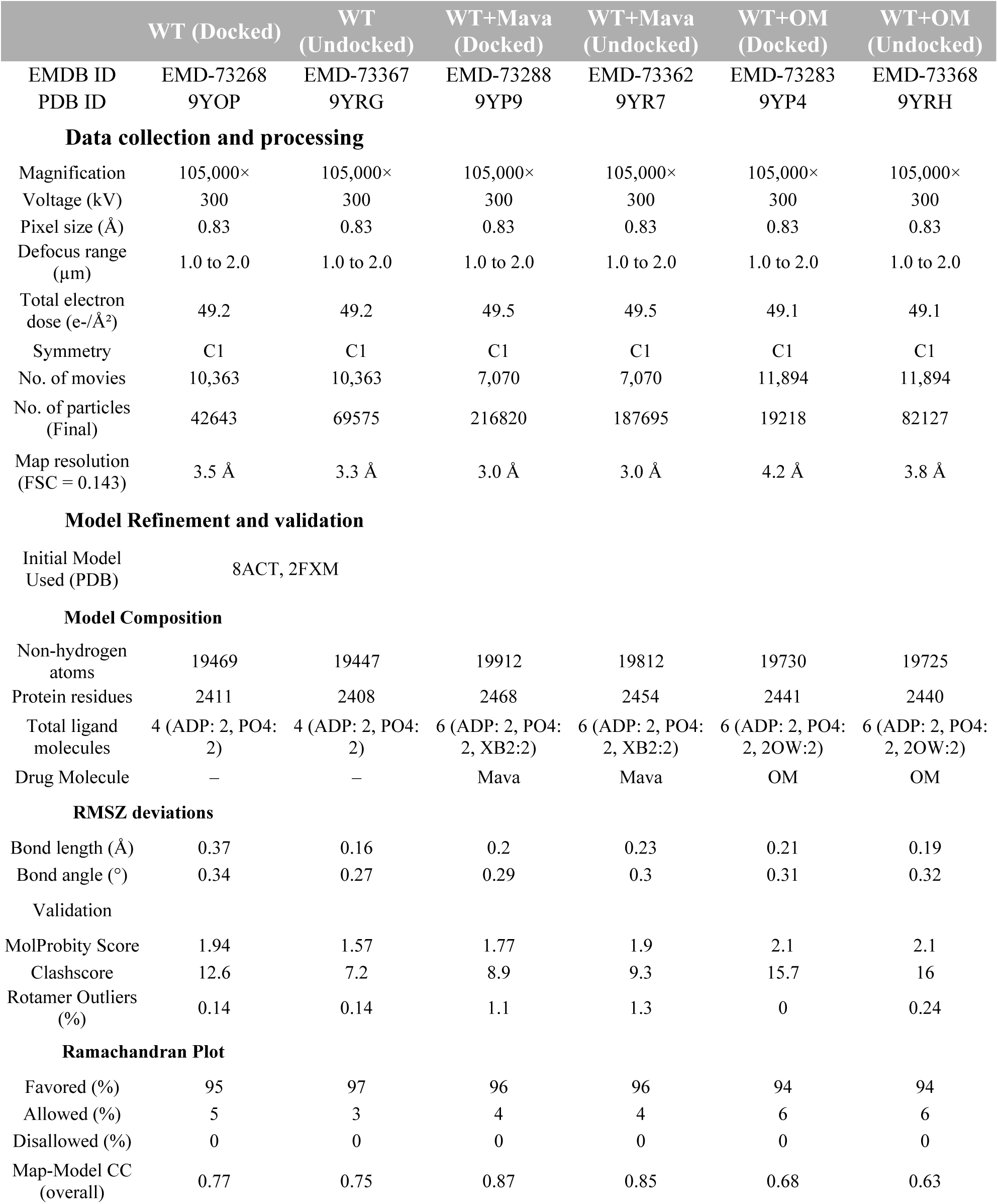
Cryo-EM data collection, refinement and validation statistics.

|  | WT (Docked) | WT (Undocked) | WT+Mava (Docked) | WT+Mava (Undocked) | WT+OM (Docked) | WT+OM (Undocked) |
| --- | --- | --- | --- | --- | --- | --- |
| EMDB ID | EMD-73268 | EMD-73367 | EMD-73288 | EMD-73362 | EMD-73283 | EMD-73368 |
| PDB ID | 9YOP | 9YRG | 9YP9 | 9YR7 | 9YP4 | 9YRH |
| Data collection and processing |  |  |  |  |  |  |
| Magnification | 105,000× | 105,000× | 105,000× | 105,000× | 105,000× | 105,000× |
| Voltage (kV) | 300 | 300 | 300 | 300 | 300 | 300 |
| Pixel size (Å) | 0.83 | 0.83 | 0.83 | 0.83 | 0.83 | 0.83 |
| Defocus range (μm) | 1.0 to 2.0 | 1.0 to 2.0 | 1.0 to 2.0 | 1.0 to 2.0 | 1.0 to 2.0 | 1.0 to 2.0 |
| Total electron dose (e-/Å²) | 49.2 | 49.2 | 49.5 | 49.5 | 49.1 | 49.1 |
| Symmetry | C1 | C1 | C1 | C1 | C1 | C1 |
| No. of movies | 10,363 | 10,363 | 7,070 | 7,070 | 11,894 | 11,894 |
| No. of particles (Final) | 42643 | 69575 | 216820 | 187695 | 19218 | 82127 |
| Map resolution (FSC = 0.143) | 3.5 Å | 3.3 Å | 3.0 Å | 3.0 Å | 4.2 Å | 3.8 Å |
| Model Refinement and validation |  |  |  |  |  |  |
| Initial Model Used (PDB) | 8ACT, 2FXM |  |  |  |  |  |
| Model Composition |  |  |  |  |  |  |
| Non-hydrogen atoms | 19469 | 19447 | 19912 | 19812 | 19730 | 19725 |
| Protein residues | 2411 | 2408 | 2468 | 2454 | 2441 | 2440 |
| Total ligand molecules | 4 (ADP: 2, PO4: 2) | 4 (ADP: 2, PO4: 2) | 6 (ADP: 2, PO4: 2, XB2:2) | 6 (ADP: 2, PO4: 2, XB2:2) | 6 (ADP: 2, PO4: 2, 2OW:2) | 6 (ADP: 2, PO4: 2, 2OW:2) |
| Drug Molecule | – | – | Mava | Mava | OM | OM |
| RMSZ deviations |  |  |  |  |  |  |
| Bond length (Å) | 0.37 | 0.16 | 0.2 | 0.23 | 0.21 | 0.19 |
| Bond angle (°) | 0.34 | 0.27 | 0.29 | 0.3 | 0.31 | 0.32 |
| Validation |  |  |  |  |  |  |
| MolProbity Score | 1.94 | 1.57 | 1.77 | 1.9 | 2.1 | 2.1 |
| Clashscore | 12.6 | 7.2 | 8.9 | 9.3 | 15.7 | 16 |
| Rotamer Outliers (%) | 0.14 | 0.14 | 1.1 | 1.3 | 0 | 0.24 |
| Ramachandran Plot |  |  |  |  |  |  |
| Favored (%) | 95 | 97 | 96 | 96 | 94 | 94 |
| Allowed (%) | 5 | 3 | 4 | 4 | 6 | 6 |
| Disallowed (%) | 0 | 0 | 0 | 0 | 0 | 0 |
| Map-Model CC (overall) | 0.77 | 0.75 | 0.87 | 0.85 | 0.68 | 0.63 |

**Table S2.** Classification of HMM particles (Open Vs Closed) based on automatic blob picking analysis. . Summary of particle distribution obtained from automated blob picking in cryoSPARC. The table lists the number of closed HMM particles corresponding to the IHM particles, and single motor domain particles, which are interpreted as open conformations. As each HMM contains two motor domains, the number of single motor domain particles was halved to estimate the total number of open HMMs. The total number of particles in each category is listed along with the percentage of closed and open HMM particles which provides an estimate of the conformational population distribution in each sample.

| Sample | Closed HMM | Single Motor Domain | Open HMM | Total HMM | Closed HMM (%) | Open HMM (%) |
| --- | --- | --- | --- | --- | --- | --- |
| WT+Mava | 301305 | 358353 | 179176 | 480481 | 62.7 | 37.3 |
| WT | 97106 | 629354 | 314677 | 411783 | 23.6 | 76.4 |
| WT+OM | 48265 | 797531 | 398765 | 447030 | 10.8 | 89.2 |

**Table S3.** Classification of HMM particles (Docked Vs Undocked) based on 3D classification.

| Sample | Docked<br>conformation | Undocked<br>conformation | Total | Docked (%) | Undocked<br>(%) |
| --- | --- | --- | --- | --- | --- |
| WT+Mava | 216820 | 187695 | 404515 | 54 | 46 |
| WT | 42643 | 69575 | 112218 | 38 | 62 |
| WT+OM | 19218 | 82127 | 101345 | 19 | 81 |

**Table S4.**
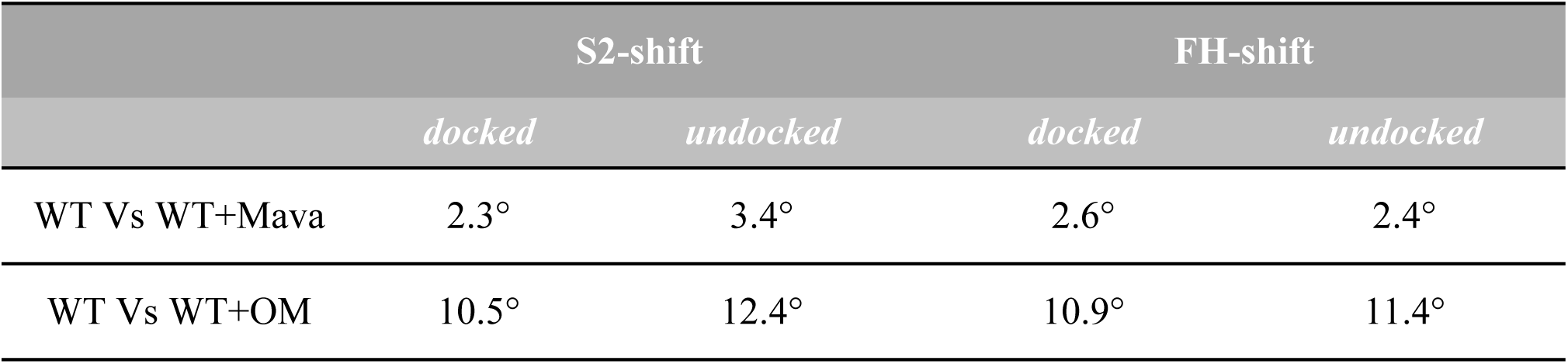
S2 and FH angular deviations in the presence of Mava and OM compared to apo WT conformations.

**Table S5.** Lever arm angular deviations in the presence of Mava and OM compared to apo WT conformations.

| Deviation measurement | Docked Conformations |  |  | Undocked Conformations |  |  |
| --- | --- | --- | --- | --- | --- | --- |
|  | *WT V<sub>S</sub>*<br>*WT+Mava* | *WT V<sub>S</sub>*<br>*WT+OM* | *WT+Mava*<br>*V<sub>S</sub>*<br>*WT+OM* | *WT V<sub>S</sub>*<br>*WT+Mava* | *WT V<sub>S</sub>*<br>*WT+OM* | *WT+Mava*<br>*V<sub>S</sub>*<br>*WT+OM* |
| BH lever arm | 1.9° | 5.6° | 5.1° | 2.3° | 4.2° | 3.5° |
| BH lever arm +<br>Proximal S2 | 1.7° | 6.2° | 5.9° | 2.0° | 4.7° | 3.9° |
| FH lever arm | 5.3° | 13.1° | 11.2° | 3.0° | 13.2° | 10.4° |
| FH lever arm +<br>Proximal S2 | 2.6° | 11.5° | 11.2° | 2.7° | 11.5° | 9.1° |

## References

1. M. A. Geeves, K. C. Holmes, Structural mechanism of muscle contraction. Annu Rev Biochem 68, 687–728 (1999).

2. J. Robert-Paganin, O. Pylypenko, C. Kikuti, H. L. Sweeney, A. Houdusse, Force Generation by Myosin Motors: A Structural Perspective. Chem Rev 120, 5–35 (2020).

3. A. Houdusse, V. N. Kalabokis, D. Himmel, A. G. Szent-Gyorgyi, C. Cohen, Atomic structure of scallop myosin subfragment S1 complexed with MgADP: a novel conformation of the myosin head. Cell 97, 459–470 (1999).

4. I. Rayment et al., Three-dimensional structure of myosin subfragment-1: a molecular motor. Science 261, 50–58 (1993).

5. J. Squire, The Structural Basis of Muscular Contraction. (Springer New York, NY, 1981).

6. D. Dutta, V. Nguyen, K. S. Campbell, R. Padron, R. Craig, Cryo-EM structure of the human cardiac myosin filament. Nature 623, 853–862 (2023).

7. R. Cooke, The role of the myosin ATPase activity in adaptive thermogenesis by skeletal muscle. Biophys Rev 3, 33–45 (2011).

8. M. A. Stewart, K. Franks-Skiba, S. Chen, R. Cooke, Myosin ATP turnover rate is a mechanism involved in thermogenesis in resting skeletal muscle fibers. Proc Natl Acad Sci U S A 107, 430–435 (2010).

9. R. Craig, R. Padron, Structural basis of the super- and hyper-relaxed states of myosin II. J Gen Physiol 154, (2022).

10. L. Alamo et al., Three-dimensional reconstruction of tarantula myosin filaments suggests how phosphorylation may regulate myosin activity. J Mol Biol 384, 780–797 (2008).

11. J. L. Woodhead et al., Atomic model of a myosin filament in the relaxed state. Nature 436, 1195–1199 (2005).

12. T. Wendt, D. Taylor, K. M. Trybus, K. Taylor, Three-dimensional image reconstruction of dephosphorylated smooth muscle heavy meromyosin reveals asymmetry in the interaction between myosin heads and placement of subfragment 2. Proc Natl Acad Sci U S A 98, 4361–4366 (2001).

13. A. Grinzato et al., Cryo-EM structure of the folded-back state of human beta-cardiac myosin. Nat Commun 14, 3166 (2023).

14. K. H. Lee et al., Interacting-heads motif has been conserved as a mechanism of myosin II inhibition since before the origin of animals. Proc Natl Acad Sci U S A 115, E1991–E2000 (2018).

15. L. Alamo, A. Pinto, G. Sulbaran, J. Mavarez, R. Padron, Lessons from a tarantula: new insights into myosin interacting-heads motif evolution and its implications on disease. Biophys Rev 10, 1465–1477 (2018).

16. H. S. Jung, S. Komatsu, M. Ikebe, R. Craig, Head-head and head-tail interaction: a general mechanism for switching off myosin II activity in cells. Mol Biol Cell 19, 3234–3242 (2008).

17. J. A. Spudich, Hypertrophic and dilated cardiomyopathy: four decades of basic research on muscle lead to potential therapeutic approaches to these devastating genetic diseases. Biophys J 106, 1236–1249 (2014).

18. R. Yotti, C. E. Seidman, J. G. Seidman, Advances in the Genetic Basis and Pathogenesis of Sarcomere Cardiomyopathies. Annu Rev Genomics Hum Genet 20, 129–153 (2019).

19. J. A. Spudich, N. Nandwani, J. Robert-Paganin, A. Houdusse, K. M. Ruppel, Reassessing the unifying hypothesis for hypercontractility caused by myosin mutations in hypertrophic cardiomyopathy. EMBO J 43, 4139–4155 (2024).

20. A. S. Adhikari et al., beta-Cardiac myosin hypertrophic cardiomyopathy mutations release sequestered heads and increase enzymatic activity. Nat Commun 10, 2685 (2019).

21. S. Nag et al., The myosin mesa and the basis of hypercontractility caused by hypertrophic cardiomyopathy mutations. Nat Struct Mol Biol 24, 525–533 (2017).

22. L. Alamo et al., Effects of myosin variants on interacting-heads motif explain distinct hypertrophic and dilated cardiomyopathy phenotypes. Elife 6, (2017).

23. S. Duno-Miranda et al., Tail length and E525K dilated cardiomyopathy mutant alter human beta-cardiac myosin super-relaxed state. J Gen Physiol 156, (2024).

24. D. V. Rasicci et al., Dilated cardiomyopathy mutation E525K in human beta-cardiac myosin stabilizes the interacting-heads motif and super-relaxed state of myosin. Elife 11, (2022).

25. L. Alamo et al., Conserved Intramolecular Interactions Maintain Myosin Interacting-Heads Motifs Explaining Tarantula Muscle Super-Relaxed State Structural Basis. J Mol Biol 428, 1142–1164 (2016).

26. S. M. Day, J. C. Tardiff, E. M. Ostap, Myosin modulators: emerging approaches for the treatment of cardiomyopathies and heart failure. J Clin Invest 132, (2022).

27. E. M. Green et al., A small-molecule inhibitor of sarcomere contractility suppresses hypertrophic cardiomyopathy in mice. Science 351, 617–621 (2016).

28. J. R. Teerlink et al., Cardiac Myosin Activation with Omecamtiv Mecarbil in Systolic Heart Failure. N Engl J Med 384, 105–116 (2021).

29. C. Y. Ho et al., Evaluation of Mavacamten in Symptomatic Patients With Nonobstructive Hypertrophic Cardiomyopathy. J Am Coll Cardiol 75, 2649–2660 (2020).

30. T. Kampourakis, X. Zhang, Y. B. Sun, M. Irving, Omecamtiv mercabil and blebbistatin modulate cardiac contractility by perturbing the regulatory state of the myosin filament. J Physiol 596, 31–46 (2018).

31. R. L. Anderson et al., Deciphering the super relaxed state of human beta-cardiac myosin and the mode of action of mavacamten from myosin molecules to muscle fibers. Proc Natl Acad Sci U S A 115, E8143–E8152 (2018).

32. D. Auguin et al., Omecamtiv mecarbil and Mavacamten target the same myosin pocket despite opposite effects in heart contraction. Nat Commun 15, 4885 (2024).

33. S. N. McMillan, J. R. T. Pitts, B. Barua, D. A. Winkelmann, C. A. Scarff, Mavacamten inhibits myosin activity by stabilising the myosin interacting-heads motif and stalling motor force generation. bioRxiv, (2025).

34. F. H. C. Crick, The packing of α-helices: simple coiled-coils. Acta Crystallographica 6, 689–697 (1953).

35. W. Blankenfeldt, N. H. Thoma, J. S. Wray, M. Gautel, I. Schlichting, Crystal structures of human cardiac beta-myosin II S2-Delta provide insight into the functional role of the S2 subfragment. Proc Natl Acad Sci U S A 103, 17713–17717 (2006).

36. M. C. Childers, M. A. Geeves, M. Regnier, Interacting myosin head dynamics and their modification by 2’-deoxy-ADP. Biophys J 123, 3997–4008 (2024).

37. R. R. Goluguri et al., A FRET assay to monitor different structural states of human beta-cardiac myosin including the interacting-heads motif. Proc Natl Acad Sci U S A 122, e2504562122 (2025).

38. R. C. Cail, F. A. Baez-Cruz, D. A. Winkelmann, Y. E. Goldman, E. M. Ostap, Dynamics of beta-cardiac myosin between the super-relaxed and disordered-relaxed states. J Biol Chem 301, 108412 (2025).

39. N. Nandwani et al., Hypertrophic cardiomyopathy mutations Y115H and E497D disrupt the folded-back state of human beta-cardiac myosin allosterically. Nat Commun 16, 8751 (2025).

40. D. Dutta et al., Thick filament molecular interfaces play a critical role in pathogenesis of hypertrophic and dilated cardiomyopathy. bioRxiv, (2025).

41. J. A. Rohde, O. Roopnarine, D. D. Thomas, J. M. Muretta, Mavacamten stabilizes an autoinhibited state of two-headed cardiac myosin. Proc Natl Acad Sci U S A 115, E7486–E7494 (2018).

42. M. S. Woody et al., Positive cardiac inotrope omecamtiv mecarbil activates muscle despite suppressing the myosin working stroke. Nat Commun 9, 3838 (2018).

43. V. J. Planelles-Herrero, J. J. Hartman, J. Robert-Paganin, F. I. Malik, A. Houdusse, Mechanistic and structural basis for activation of cardiac myosin force production by omecamtiv mecarbil. Nat Commun 8, 190 (2017).

44. D. A. Winkelmann, E. Forgacs, M. T. Miller, A. M. Stock, Structural basis for drug-induced allosteric changes to human beta-cardiac myosin motor activity. Nat Commun 6, 7974 (2015).

45. R. Sharma, J. Ge, C. M. Yengo, R. Craig, R. Padron, BPS2025 - Cardiac myosin 25-hep E525K mutated construct with mavacamten is stabilized by three blocked head subfragment-2 interactions. Biophysical Journal 124, (2025).

46. Z. Ujfalusi et al., Dilated cardiomyopathy myosin mutants have reduced force-generating capacity. J Biol Chem 293, 9017–9029 (2018).

47. M. Schorb, I. Haberbosch, W. J. H. Hagen, Y. Schwab, D. N. Mastronarde, Software tools for automated transmission electron microscopy. Nat Methods 16, 471–477 (2019).

48. A. Punjani, J. L. Rubinstein, D. J. Fleet, M. A. Brubaker, cryoSPARC: algorithms for rapid unsupervised cryo-EM structure determination. Nat Methods 14, 290–296 (2017).

49. T. Bepler et al., Positive-unlabeled convolutional neural networks for particle picking in cryo-electron micrographs. Nat Methods 16, 1153–1160 (2019).

50. E. C. Meng et al., UCSF ChimeraX: Tools for structure building and analysis. Protein Sci 32, e4792 (2023).

51. R. T. Kidmose et al., Namdinator - automatic molecular dynamics flexible fitting of structural models into cryo-EM and crystallography experimental maps. IUCrJ 6, 526–531 (2019).

52. P. V. Afonine et al., Real-space refinement in PHENIX for cryo-EM and crystallography. Acta Crystallogr D Struct Biol 74, 531–544 (2018).

53. A. Casanal, B. Lohkamp, P. Emsley, Current developments in Coot for macromolecular model building of Electron Cryo-microscopy and Crystallographic Data. Protein Sci 29, 1069–1078 (2020).

54. C. J. Williams et al., MolProbity: More and better reference data for improved all-atom structure validation. Protein Sci 27, 293–315 (2018).

